# Stimulation of the catalytic activity of the tyrosine kinase Btk by the adaptor protein Grb2

**DOI:** 10.1101/2022.08.25.505243

**Authors:** Laura M. Nocka, Jay T. Groves, John Kuriyan

## Abstract

The Tec-family kinase Btk contains a lipid-binding Pleckstrin homology and Tec homology (PH-TH) module connected by a proline-rich linker to a “Src module”, an SH2-SH3-kinase unit also found in Src-family kinases and Abl. We showed previously that Btk is activated by PH-TH dimerization, which is triggered on membranes by the phosphatidyl inositol phosphate PIP_3_, or in solution by hexakisinositol phosphate (IP_6_) (Wang *et al.* 2015, https://doi.org/10.7554/eLife.06074). We now report that the ubiquitous adaptor protein growth-factor-receptor-bound protein 2 (Grb2) binds to and substantially increases the activity of PIP_3_-bound Btk on membranes. Using reconstitution on supported-lipid bilayers, we find that Grb2 can be recruited to membrane-bound Btk through interaction with the proline-rich linker in Btk. This interaction requires intact Grb2, containing both SH3 domains and the SH2 domain, but does not require that the SH2 domain be able to bind phosphorylated tyrosine residues – thus Grb2 bound to Btk is free to interact with scaffold proteins via the SH2 domain. We show that the Grb2-Btk interaction recruits Btk to scaffold-mediated signaling clusters in reconstituted membranes. Our findings indicate that PIP_3_-mediated dimerization of Btk does not fully activate Btk, and that Btk adopts an autoinhibited state at the membrane that is released by Grb2.

## Introduction

B cell signaling relies on the sequential activation of three tyrosine kinases that transduce the stimulation of the B cell receptor (BCR) into the generation of calcium flux and the initiation of signaling pathways, such as the Ras/MAPK pathway (Engels, Engelke, & Wienands, 2008; Weiss & Littman, 1994). Activation of a Src-family kinase, Lyn, results in the phosphorylation of ITAMs (Immunoreceptor Tyrosine-based Activation Motifs) in the cytoplasmic tails of the B cell receptor. This results in the recruitment and activation of the tyrosine kinase Syk, which phosphorylates the scaffold protein SLP65 (SH2 domain-containing Leukocyte Protein of 65 kDa) also known as BLNK (B-cell Linker Protein) (Kurosaki & Tsukada, 2000). The activation of Syk triggers the recruitment of diverse signaling proteins to the plasma membrane, including the Tec-family kinase Btk, which is critical for the development and proliferation of mature B cells (Figure 1A) (Hendriks, Yuvaraj, & Kil, 2014; Rip, Van Der Ploeg, Hendriks, & Corneth, 2018; Scharenberg, Humphries, & Rawlings, 2007; Tsukada et al., 1993). Signaling by the T cell receptor follows a similar pathway, with the Src-family kinase Lck activating the Syk-family kinase Zap-70 (Zeta-chain-associated protein kinase of 70 kDa). Zap-70 phosphorylates the scaffold protein LAT (Linker for Activation of T cells), which leads to the activation of the Tec-family kinase Itk (Interleukin-2-inducible T cell Kinase) (Courtney, Lo, & Weiss, 2018; Weiss & Littman, 1994).

**Figure 1.**
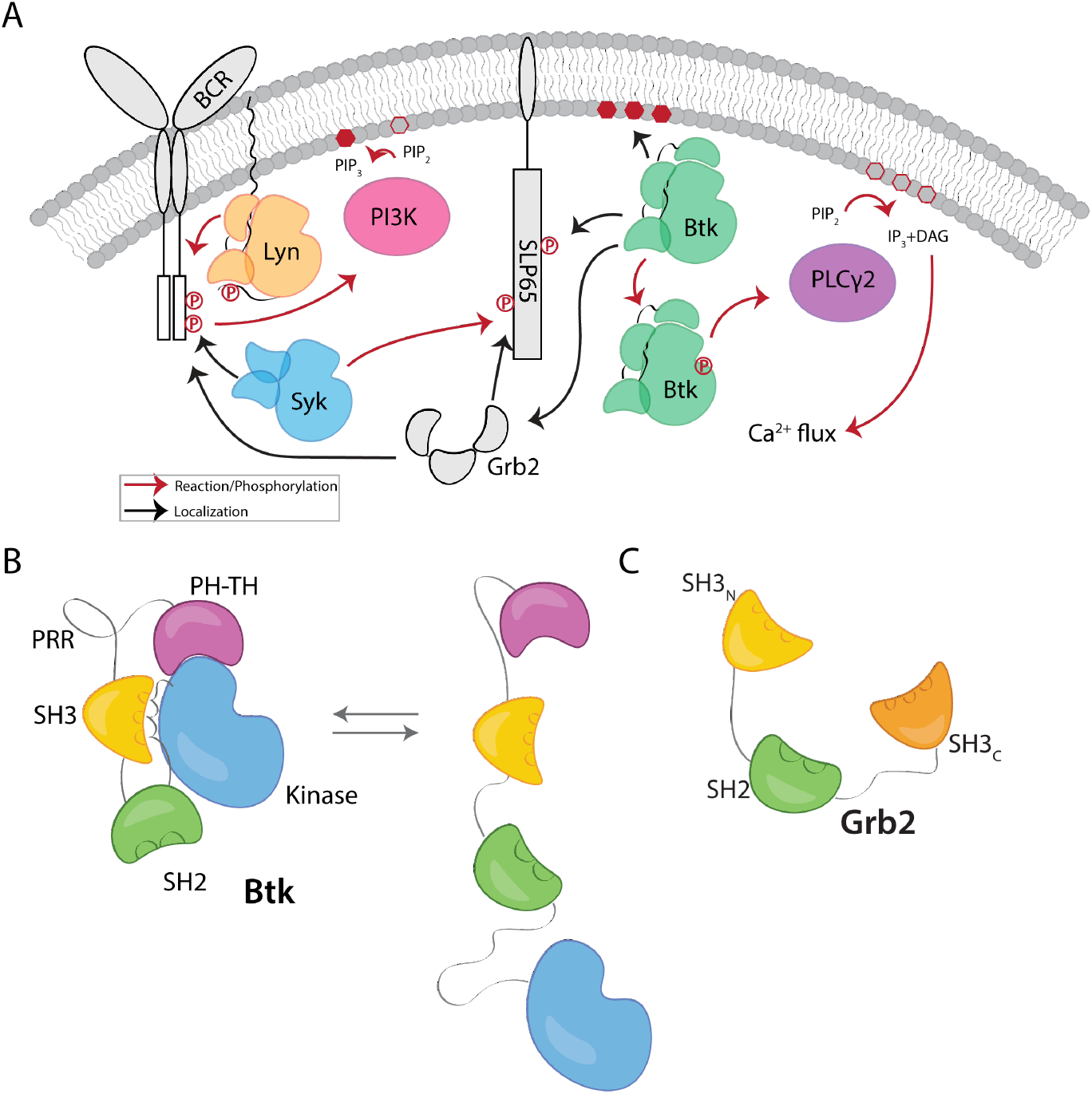
Btk and Grb2 are integral to B-cell signaling. A) The B cell signaling cascade relies on kinases and adaptor molecules. B) Btk is made up of five distinct domains. The Pleckstrin homology and Tec homology domains fold together into one module (PH-TH), which is connected through a linker containing two proline-rich regions (PRR) to the Src module (SH3, SH2 and kinase domain). These domains together mediate autoinhibition of Btk. C) Grb2 domain architecture consists of two SH3 domains that flank an SH2 domain.

Btk has an N-terminal PH-TH module, consisting of a Pleckstrin-homology (PH) domain followed by a Zinc-bound Tec-homology (TH) domain. The PH-TH module is connected through a proline-rich linker to a Src module, comprising an SH3 domain, an SH2 domain, and a kinase domain (Figure 1B) (Joseph, Wales, Fulton, Engen, & Andreotti, 2017; Shah, Amacher, Nocka, & Kuriyan, 2018; Wang et al., 2015). The Src module of Btk is structurally similar to the corresponding modules of other cytoplasmic tyrosine kinases such as c-Src, Lck and Abl. SH2 domains bind to phosphotyrosine-containing peptides, while SH3 domains recognize specific PxxP motifs within proline-rich peptide segments. In the autoinhibited forms of kinases with Src modules, the SH3 domain and the N-terminal lobe of the kinase domain sandwich the SH2-kinase linker, and the SH2 domain sits adjacent to the C-terminal lobe of the kinase domain. Together these contacts prevent the kinase domain from adopting an active conformation (Amatya, Lin, & Andreotti, 2019; Shah et al., 2018). The PH-TH module of Btk is connected to the Src module by a 44-residue linker that contains ten proline residues and two PxxP motifs that are potential SH3-binding sites. Btk is recruited to the membrane through binding of PIP_3_ by the PH-TH module (Baraldi et al., 1999; Chung et al., 2019; Hyvönen & Saraste, 1997).

Crystal structures of the Btk PH-TH module, first determined by the late Matte Saraste and colleagues, reveal a dimeric arrangement, which we refer to as the “Saraste dimer” (Baraldi et al., 1999; Hyvönen & Saraste, 1997). Mutagenesis of residues at the Saraste-dimer interface has shown that the PH-TH dimer is critical for Btk activity in cells (Chung et al., 2019). Btk is activated *in vitro* by vesicles containing PIP_3_, or by inositol hexakisphosphate (IP_6_) in solution (Kim et al., 2019; Wang et al., 2015). The activation by IP_6_ requires formation of the Saraste dimer, and involves the binding of two IP_6_ molecules, one bound at the canonical PIP-binding site in the PH domain and the other to a peripheral site. The physiological relevance of the IP_6_-stimulated Btk activity in T cell-independent B cell activation has been demonstrated recently (Kim et al., 2019). Experiments on supported-lipid bilayers, as well as molecular dynamics simulations, have shown that PIP_3_ in membranes promotes dimerization of the PH-TH module through binding to the canonical and peripheral sites (Chung et al., 2019; Wang, Pechersky, Sagawa, Pan, & Shaw, 2019).

Our understanding of the structure of Btk is based on structures determined separately for the isolated PH-TH module, the Src module, and a construct in which the PH-TH module is linked artificially to the kinase domain (Baraldi et al., 1999; Hyvönen & Saraste, 1997; Wang et al., 2015). All four domains of Btk participate in maintaining the autoinhibited state (Figure 1B). Deletion of the PH-TH domain from full-length Btk results in a more active kinase *in vitro* (Wang et al., 2015). NMR analysis demonstrates that there is an autoinhibited form of Btk in which the PH-TH module interacts with the kinase domain differently than observed in the crystal structure of the PH-TH-kinase fusion, but the structure of this state remain to be defined in detail (Joseph et al., 2017). Additionally through mutational studies and SAXS measurements an SH2-kinase interface that stabilizes an active form of the Btk has been identified (Duarte et al., 2020). Together these studies suggest a dynamic equilibrium between active and autoinhibited forms of the kinase.

Adaptor proteins play critical roles in facilitating the organization of enzymes during signal transduction (Hashimoto et al., 1999; Koretzky, Abtahian, & Silverman, 2006; Rodriguez et al., 2001). Growth-factor-receptor-bound protein 2 (Grb2) is a ubiquitous adaptor protein that is important for receptor tyrosine kinase signaling and the mitogen-activated protein kinase (MAPK) pathway (Cantor, Shah, & Kuriyan, 2018; Clark, Stern, & Horvitz, 1992; Gale, Kaplan, Lowenstein, Schlessinger, & Bar‑Sagi, 1993; Lowenstein et al., 1992; Olivier et al., 1993). Grb2 consists of two SH3 domains that flank an SH2 domain (Figure 1C). Grb2 is responsible for bringing signaling enzymes to their substrates and also helps in the formation of signaling clusters at the plasma membrane by virtue of its ability to crosslink scaffold proteins or receptors (Huang et al., 2019; Huang, Chiang, & Groves, 2017; Huang et al., 2016; C.‑W. Lin et al., 2022; Su et al., 2016). For example, in the MAPK pathway, the SH3 domains of two Grb2 molecules bind to the Ras activator Son of Sevenless (SOS). Grb2 binds to scaffold proteins, such as SLP65 in B cells and LAT in T cells, through interaction of the Grb2 SH2 domain with phosphotyrosine residues on the scaffold proteins (Engels et al., 2009; Reif, Buday, Downward, & Cantrell, 1994). In this way, Grb2 recruits SOS to the scaffold proteins or receptors, and the interaction of one SOS molecule with two Grb2 molecules promotes crosslinking of these proteins. In T cells, a protein closely related to Grb2, GADS (Grb2-related adaptor downstream of Shc), works together with Grb2 to crosslink scaffolding proteins and enzymes downstream of the TCR (T cell receptor) (Liu, Fang, Koretzky, & McGlade, 1999). Grb2 is also reported to be capable of dimerization, and dimeric Grb2 hinders basal signaling of the fibroblast growth factor receptor 2 (FGFR2), thereby tuning the activity of the receptor (C.‑C. Lin et al., 2012).

Memory-type B cells expressing membrane-bound IgG (mIgG) as a component of the BCR rely on an immunoglobulin tail tyrosine (ITT) motif that is phosphorylated by Syk and recruits Grb2 through binding of its SH2 domain to the resulting phosphotyrosine residues. It was shown, by fusing domains of Grb2 to the IgG tail (in the absence of the ITT motif), that the N-terminal SH3 domain of Grb2 plays a critical role in generating downstream signals. Affinity purifications using the isolated N-terminal SH3 domain of Grb2 showed that this domain interacts with Btk, as well as with the ubiquitin ligase Cbl and SOS (Engels et al., 2014).

Substitution of the N-terminal SH3 domain of Grb2 by the C-terminal SH3 domain of Grb2, or by the N-terminal SH3 domain of the Grb2 family member GRAP resulted in failure to trigger Ca^2+^ flux. In the mIgG fusion with the N-terminal SH3 domain of GRAP, Ca^2+^ flux was restored by the substitution of three residues in the GRAP SH3 domain by the corresponding residues in Grb2 (Engels et al., 2014). Thus, an interaction between Grb2 and Btk plays a critical role in BCR signaling.

We now report the discovery, made using *in-vitro* reconstitution of Btk on PIP_3_-containing membranes, that Btk and Grb2 interact through the proline-rich linker of Btk, and that, through this interaction, Grb2 strongly stimulates the kinase activity of Btk. Our data indicate that membrane recruitment of Btk by PIP_3_ is insufficient for potent activation of Btk, which requires interaction with Grb2 at the membrane. The activation of Btk by Grb2 relies primarily on the N-terminal SH3 domain of Grb2 and does not require that the SH2 domain be able to bind to phosphotyrosine residues – we infer that the SH2 domain is free to interact with phosphotyrosine residues presented by scaffold proteins. Our data suggest a potential mechanism whereby the binding of one SH3 domain of Grb2 to the proline-rich linker of Btk enables the other SH3 domain to bind to the SH2-kinase linker of Btk, thereby displacing the inhibitory intramolecular interaction of the Btk SH3 domain. Thus, the interaction between Grb2 and Btk can integrate the localization of Btk with its activation.

## Results and Discussion

### Btk recruits Grb2 to PIP_3_-containing supported-lipid bilayers through the proline-rich region of Btk

To probe the interaction between Btk and Grb2, we utilized a supported-lipid bilayer system that we had used previously to characterize the interaction of the isolated Btk PH-TH module with lipids (Chung et al., 2019). Using this system we discovered, unexpectedly, that Grb2 can be recruited to the membrane via an interaction with membrane-bound full-length Btk. We had shown previously that the PH-TH module of Btk is recruited to membranes containing 4% PIP_3_, and that this recruitment exhibits a sharp dependence on PIP_3_ concentration in the membrane. To characterize the interaction between Grb2 and full-length Btk we used supported-lipid bilayers containing 4% PIP_3_, to which full-length Btk is recruited from solution via the PH-TH module (Chung et al., 2019). We found that recruitment of Btk to the membrane also resulted in Grb2 being recruited to the membrane.

Recruitment of Grb2 to the membrane was measured using total internal reflection fluorescence (TIRF) microscopy and Grb2 labeled with Alexa Fluor 647 (Grb2-647). The surface sensitivity of TIRF imaging provides a highly selective measurement of membrane-associated Grb2-647, without picking up signal from protein in the solution phase (Huang, Ditlev, Chiang, Rosen, & Groves, 2017). When Btk and Grb2-647 were added together, Grb2-647 was recruited to the bilayer, as indicated by an increase in fluorescence (Figure 2A). When Grb2-647 was added to the supported-lipid bilayers without Btk there was no change in fluorescence above background (Figure 2A). These experiments indicate that Grb2 is capable of binding directly to Btk in the absence of other proteins. We confirmed that the observed fluorescence change was not due to an irreversible process, such as protein aggregation, by showing that the increase in fluorescence could be reversed rapidly by addition of unlabeled Grb2 at a five-fold higher solution concentration than labeled Grb2 (Figure 2B).

**Figure 2.**
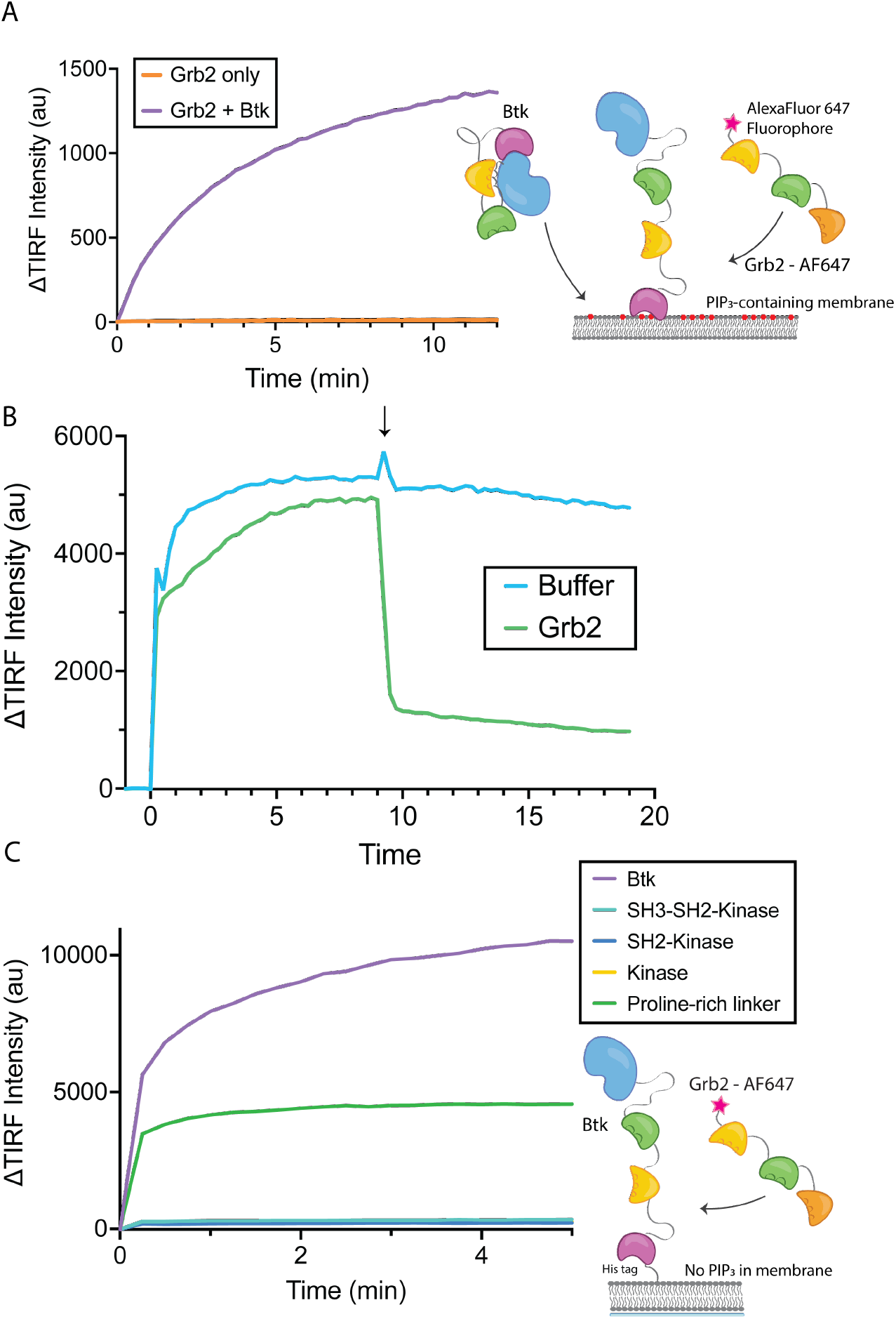
Grb2 interacts with membrane-bound Btk. A) Fluorescently labeled Grb2 was added to supported-lipid bilayers containing 4% PIP_3_, with or without Btk. The change in fluorescence intensity, monitored by TIRF, is plotted over time. B) The binding of Grb2 to membrane bound Btk is reversible. Grb2 was added to a bilayer decorated with His-tagged Btk. In the blue curve, the bilayer is washed with buffer, the arrow indicates when the wash occurred. In one experiment (blue graph), the bilayer was washed with buffer. In another experiment (green graph), the bilayer was washed with a solution containing 10-fold higher concentration of unlabeled Grb2. C) Various constructs of Btk (Btk, SH3-SH2-Kinase, SH2-Kinase, Proline-rich linker) with His tags were tethered to supported-lipid bilayers containing 4% DGS-NTA(Ni) lipids. The bilayers were washed after equilibration to remove any weakly bound Btk. Fluorescently-labeled Grb2 was added to these bilayers and the change in fluorescence intensity with time is shown. Each experiment shown here was carried out in at least triplicate, though only one experiment is plotted for each as exact intensity varies with each bilayer preparation.

We next addressed the question of which regions of Btk are necessary for the interaction with Grb2. To do this, we tethered various constructs of Btk to membranes containing DGS-NTA(Ni) (1,2-dioleoyl-sn-glycero-3-[(N-(5-amino-1-carboxypentyl)iminodiacetic acid)succinyl] (nickel salt)) lipids by using an N-terminal hexa-histidine tag on Btk, rather than relying on the binding of the PH-TH module to PIP_3_. Constructs of N-terminally His-tagged Btk could then be tethered directly to the membrane through the binding of the histidine tag to DGS-NTA(Ni) lipids. This tethering method has been shown to result in limited unbinding of the His-tagged protein over the timescale of our experiments (less than one hour) (Nye & Groves, 2008). This allowed us to study the binding of Grb2 to constructs of Btk that do not contain the PH-TH module (see Materials and Methods for precise definition of the Btk and Grb2 constructs).

Supported-lipid bilayers containing 4% DGS-NTA(Ni) were prepared and His-tagged Btk constructs were added to these bilayers at different concentrations. Each of the following constructs was tested by adding Grb2-647 and monitoring the change in TIRF intensity at the membrane: full-length Btk, Btk in which the PH-TH module and proline-rich linker are deleted (SH3-SH2-kinase; residues 212-659 of human Btk), SH2-kinase (residues 281-659), the kinase domain alone (residues 402-659), and the proline-rich linker alone (residues 171-214). Grb2-647 is recruited to the bilayer when full-length Btk or the isolated proline-rich linker are tethered to the membrane. The tethering of other constructs to the membrane did not show an intensity change above background upon addition of labeled Grb2 (Figure 2C).

The Btk constructs used in these experiments were expressed using a bacterial system in which co-expression of a tyrosine phosphatase is expected to maintain the proteins in the unphosphorylated state, despite the presence of the Btk kinase domain in some of the constructs (Seeliger et al., 2005; Wang et al., 2015). We checked the phosphorylation state of full-length Btk used in these experiments by Western blot analysis of the protein with a pan-phosphotyrosine antibody, as well as by mass spectrometry, and observed that purified Btk was not phosphorylated. The interaction of Btk with Grb2 is therefore not dependent on an SH2-phosphotyrosine interaction.

These experiments demonstrate that the proline-rich linker of Btk is able to recruit Grb2 to the membrane, without the other domains of Btk being present. No binding is detected to constructs which lack the PH-TH module and the proline-rich linker. The set of experiments with His-tagged Btk also show that the interaction between Btk and Grb2 is not dependent on Btk binding to PIP_3_ in the membrane, since there is no PIP_3_ present in these experiments. We conclude that the proline-rich linker is a principal determinant of the interaction between Btk and Grb2.

### Grb2 enhances the kinase activity of Btk

Btk is activated by PIP_3_-containing vesicles, as shown by experiments in which the phosphorylation of full-length Btk was monitored by Western blot with pan-phosphotyrosine antibody (Wang et al., 2015). We repeated those experiments by incubating Btk (1 μM bulk solution concentration) in the presence or absence of lipid vesicles containing 4% PIP_3_, and added increasing concentrations of Grb2, from 0-10 μM bulk solution concentration. In the absence of vesicles, no change in phosphorylation is detected when Grb2 is added to Btk (Figure 3A-B, and Figure 3 – supplement 1). In the presence of PIP_3_-containing vesicles, the addition of Grb2 results in substantially increased levels of Btk phosphorylation, compared to the presence of PIP_3_-containing vesicles alone. When Btk is mixed with Grb2, the phosphorylation level detected at 5 minutes after initiation of the reaction is comparable to that seen at 20 minutes for Btk without Grb2 (Figure 3A-B and Figure 3 – supplement 1). Initiation of the kinase reaction also results in phosphorylation of Grb2 (Figure 3C and Figure 3 – supplement 2).

**Figure 3.**
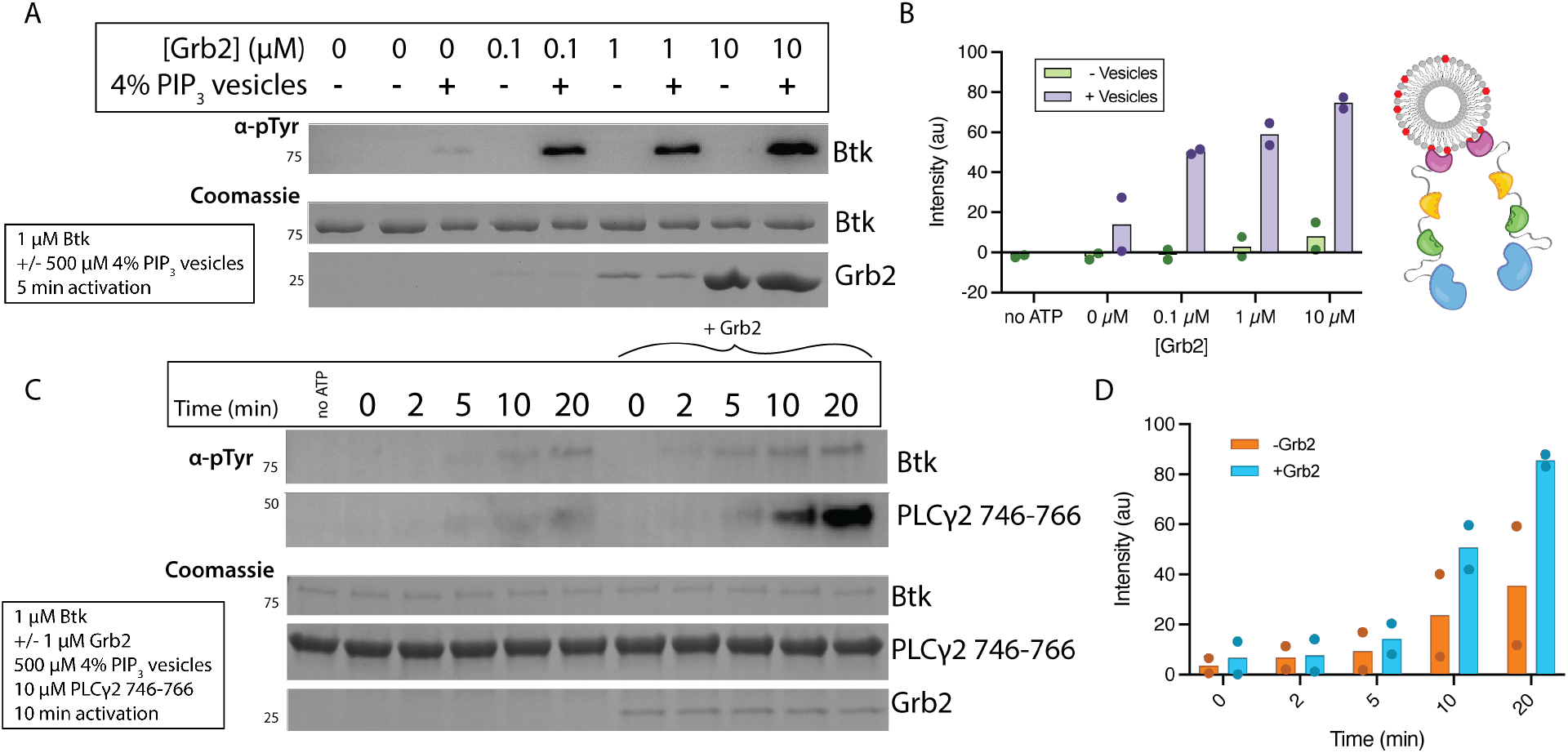
Grb2 enhances Btk kinase activity in a PIP_3_ dependent manner. The activity of Btk is monitored by measuring the phosphorylation of Btk, Grb2, and a PLCγ2-peptide fusion protein. A) Grb2 was titrated into samples containing Btk with or without 4% PIP_3_ lipid vesicles. These samples were activated for 5 minutes and then quenched. Total phosphorylation was measured by Western blot. See Figure 3 Supplement 1 for independent experiment replicate. B) Quantification of Btk phosphorylation measured from (A) along with one independent experiment replicate, bar represents the mean intensity. C) All samples contain Btk, PLCγ2-peptide fusion and 4% PIP_3_ lipid vesicles. Samples were activated and quenched at the timepoints listed (0-20 minutes), the left and right sides of the blot compare samples without or with Grb2. See Figure 3 Supplement 2 for independent experiment replicate. D) Quantification of PLCγ2-peptide fusion phosphorylation measured from (C), along with one independent experiment replicate, bar represents the mean intensity.

We tested whether the binding of Grb2 to Btk influences the ability of Btk to phosphorylate its specific substrate, PLCγ2. To do this we monitored phosphorylation of a peptide segment spanning residues 746 to 766 of PLCγ2 that contains two tyrosine residues. Phosphorylation of this segment by Btk plays a key role in activation of PLCγ2 in B cells (Ozdener, Dangelmaier, Ashby, Kunapuli, & Daniel, 2002; Rodriguez et al., 2001). This peptide segment was fused to an N-terminal SUMO protein and a C-terminal Green fluorescent protein (GFP) (referred to as the PLCγ2-peptide fusion) to allow for visualization on a gel, as the substrate is otherwise too small to analyze by Western blot. We added Btk to PIP_3_-containing vesicles in the presence of PLCγ2-peptide fusion, with or without the addition of Grb2 (Wang et al., 2015). Phosphorylation of PLCγ2-peptide fusion was measured by Western blot analysis of total phosphotyrosine. Under these conditions, the presence of Grb2 enhances phosphorylation of the PLCγ2-peptide fusion substantially (Figure 3C-D and Figure 3 – supplement 2). For example, 20 minutes after initiation of the kinase reaction, the level of phosphorylation as detected by Western blot is increased by about three times in the presence of Grb2.

Detectable activation of Btk by Grb2 only occurs when the PH-TH module of Btk engages PIP_3_ at the membrane. This is an important point, because Grb2 is a ubiquitous protein that is involved in many different signaling pathways and is expressed constitutively (Hein et al., 2015; Shi et al., 2016). Our results show that Grb2 activation of Btk is layered upon a necessary first step of PIP_3_ generation, which requires BCR stimulation. Enhanced activation of Btk results in increased phosphorylation of Btk itself as well as phosphorylation of the PLCγ2-peptide fusion.

### All three domains of Grb2 are necessary for stimulation of Btk kinase activity

We made constructs corresponding to each individual domain of Grb2 (N-terminal SH3, SH2, and C-terminal SH3), or combinations of domains (N-terminal SH3-SH2, SH2-C-terminal SH3, and N-terminal SH3-C-terminal SH3, see Materials and Methods for the specification of these constructs). We made a Grb2 variant, R86K in which a conserved arginine residue in the SH2 domain that is critical for phosphotyrosine binding is mutated to lysine. Substitution of the corresponding arginine residue in other SH2 domains attenuates the binding of the SH2 domains to phosphorylated peptides (Mayer, Jackson, Van Etten, & Baltimore, 1992). We have demonstrated recently that the R86K mutation impairs the ability of Grb2 to promote phase separation of scaffold proteins (C.‑W. Lin et al., 2022). Another Grb2 variant (Y160E) has a reduced capacity for dimerization (Ahmed et al., 2015). The ability of each of these constructs to stimulate Btk activity was tested, as measured by phosphorylation of the PLCγ2-peptide fusion (Figure 4A and Figure 4 – supplement 1). All reactions were carried out in the presence of unilamellar vesicles at a total lipid concentration of 500 μM containing 4% PIP_3_. Protein and vesicles were incubated together for 15 minutes in the absence of ATP, and the reaction time is measured from when ATP was added to the solution.

**Figure 4.**
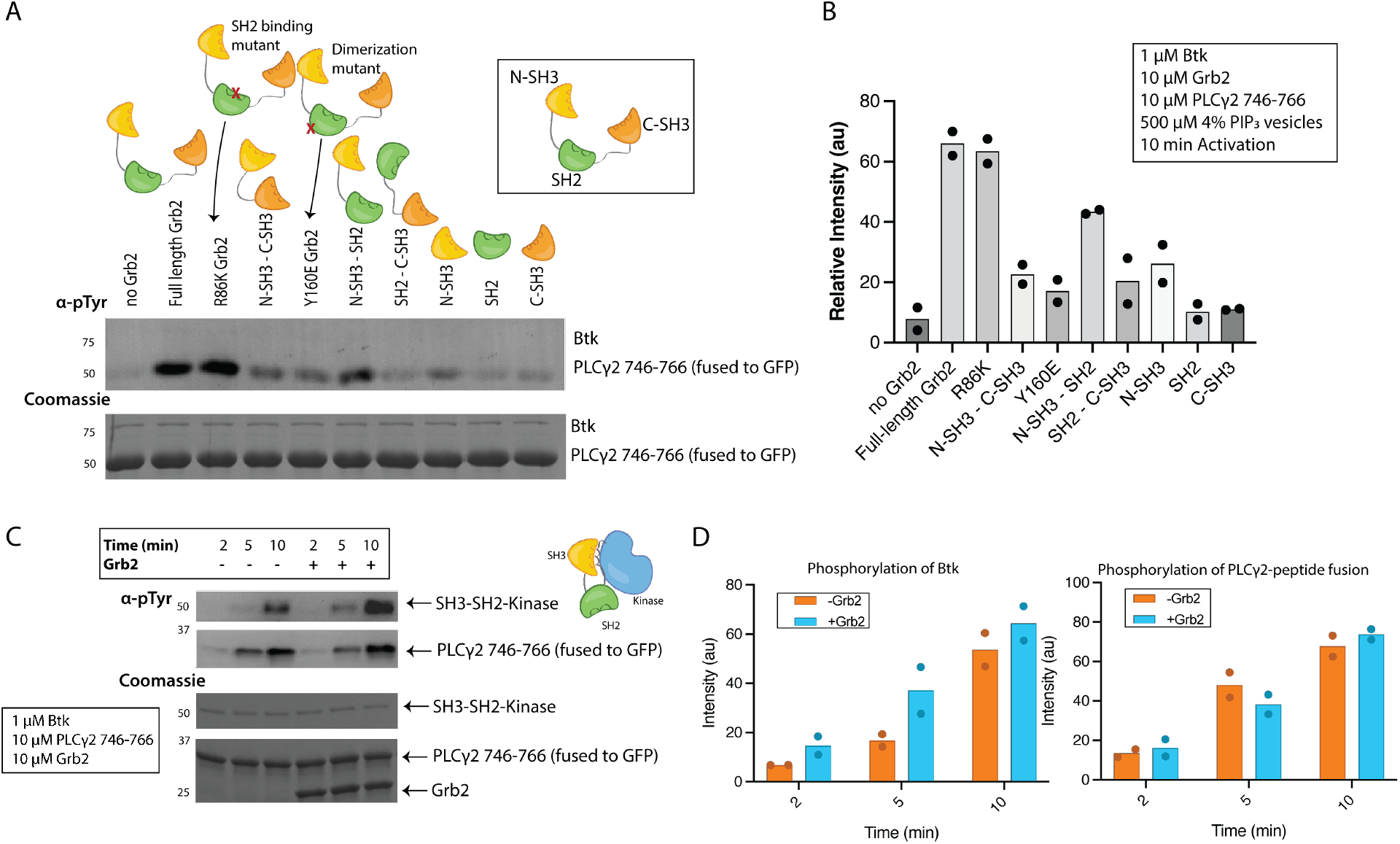
Activation of full-length and truncated Btk by different Grb2 constructs. A) Phosphorylation of PLCγ2 peptide fusion by Btk in the presence of 4% PIP_3_ lipid vesicles was monitored in the presence of various Grb2 constructs. These constructs include: full length Grb2, R86K, N-terminal SH3 fused to C-terminal SH3, Y160E, N-terminal SH3-SH2, SH2-C-terminal SH3, N-terminal SH3, SH2, and C-terminal SH3. See Figure 4 Supplement 1 for independent experiment replicate. B) Quantification of PLCγ2-peptide fusion phosphorylation measured from (A), along with one independent experiment replicate, bar represents the mean intensity. C) Activation of a truncated construct of Btk, SH3-SH2-kinase. Total phosphorylation was monitored in the absence or presence of Grb2 over time. See Figure 4 Supplement 2 for independent experiment replicate. D) Quantification of Btk phosphorylation or PLCγ2-peptide fusion phosphorylation measured from (C), along with one independent experiment replicate, bar represents the mean intensity.

Full-length Grb2 increases the phosphorylation of the PLCγ2-peptide fusion by six-fold relative to the reaction in which no Grb2 was added (Figure 4 and Figure 4 – supplement 1 and 2). The Grb2 variant in which the ability of the SH2 domain to bind to phosphopeptides is impaired (Grb2 R86K) stimulates the reaction to essentially the same extent as wild-type Grb2, indicating that the phosphopeptide-binding ability of the Grb2 SH2 domain is not required for stimulation of Btk activity by Grb2. This observation suggests that Grb2 bound to Btk will retain the capacity to dock on phosphorylated scaffold proteins via the SH2 domain, as discussed below. Removal of any of the three component domains of Grb2 results in substantial reduction of phosphorylation of the PLCγ2-peptide fusion (Figure 4 and Figure 4 – supplement 1 and 2).

The crystal structure of Grb2 (PDB code 1GRI) shows the formation of a Grb2 dimer in which there are extensive interactions between the SH3 domains of one Grb2 molecule and the SH2 domain of the other (Maignan et al., 1995). In our experiments, the stimulatory effects of Grb2 on Btk activity are observed only when Btk is localized to the membrane, and is at high local concentration, thus potentially promoting Grb2 dimerization. Deletion of the SH2 domain would disrupt the dimeric arrangement seen in the crystal structure. Mutation of Tyr 160 in Grb2, located at the distal surface of the SH2 domain and involved in the inter-subunit contacts in the crystallographic dimer, has been shown previously to affect Grb2 dimerization (Ahmed et al., 2015; C.‑C. Lin et al., 2012). More recently, we have shown that the Y160E Grb2 mutant is defective in promoting phase separation of scaffold proteins upon phosphorylation (C.‑W. Lin et al., 2021). In the present study, the use of Grb2-Y160E shows a reduction in the PLCγ2-peptide fusion phosphorylation by Btk, compared to the wild-type Grb2. This result indicates that Grb2 dimerization may be important for stimulation of Btk activity, although a definitive analysis of the mechanism awaits further study of additional Grb2 mutants.

Constructs containing only the N-terminal SH3 domain of Grb2 consistently have a larger effect on phosphorylation than constructs containing only the C-terminal SH3 domain (Figure 4A-B and Figure 4 – supplement 1). This indicates that the N-terminal SH3 domain is more important for the interaction with Grb2, consistent with the earlier finding that fusion of the N-terminal SH3 domain of Grb2 to mIgG is sufficient for promoting Ca^2+^ flux through Btk (Engels et al., 2014).

We studied a shorter Btk construct corresponding to the Src module (SH3-SH2-Kinase), which lacks the PH-TH module and the PH-TH-SH3 linker. We monitored the phosphorylation of this construct using the Western blot assay. This construct is much more active than the full-length construct and its activity is not dependent on lipids, due to the lack of a PH-TH module (Wang et al., 2015). Each reaction condition contained 1 μM Btk, 10 μM PLCγ2-peptide fusion and either 0 or 10 μM Grb2. Reactions were quenched at a range of timepoints from 2 to 10 minutes, and phosphorylation levels on Btk and PLCγ2-peptide fusion were measured separately (Fig 4C-D and Figure 4 – supplement 2). Phosphorylation on Btk increase slightly in the presence of Grb2, while the levels of phosphorylation on the PLCγ2-peptide fusion are similar with or without Grb2.

The slight stimulation by Grb2 of the autophosphorylation of the Src module construct of Btk shows that Grb2 stimulation of Btk activity may also rely on elements additional to the proline-rich linker and internal to the Src module. This points to the presence of another binding site within Btk that is capable of interacting with Grb2. The Src-module construct of Btk contains the SH2-kinase linker, which binds to the Btk SH3 domain in an intramolecular interaction in the autoinhibited state (Moarefi et al., 1997; Moroco et al., 2014). One possible explanation for the change in phosphorylation observed here is that Grb2 has some affinity for the SH2-kinase linker and is able to release autoinhibition of the kinase by competing with the Btk SH3 domain. We expect the intermolecular binding of an SH3 domain to the SH2-kinase linker of Btk to be very weak, as demonstrated by the lack of Grb2 recruitment by this construct when tethered to supported-lipid bilayers (Figure 2C). In the case of full-length Btk, this competition could be assisted by one SH3 domain of Grb2 binding to the proline-rich linker and promoting a pseudo-intramolecular binding of the second SH3 domain of Grb2 to the SH2-kinase linker of Btk. These aspects of the interaction between Btk and Grb2 await future experimentation for better understanding.

### Grb2 does not affect dimerization of Btk on the membrane and cannot rescue the catalytic activity of a dimerization-deficient mutant of Btk

Btk activation relies on homodimerization of the PH-TH modules (Chung et al., 2019; Wang et al., 2015). Given this, it is possible that the two SH3 domains of Grb2 could promote activation by facilitating dimerization by crosslinking two different Btk molecules. To check whether Grb2 impacts the dimerization of Btk we measured the diffusion constant and the dwell time of Btk on the membrane in the presence or absence of Grb2 (Figure 5A and Figure 5 – supplement 1) (Chung et al., 2019). Although there is no simple relation connecting two-dimensional diffusion and molecular complex size as there is for three-dimensional diffusion, two-dimensional diffusion on a membrane surface nonetheless changes markedly between monomers and dimers and is a sensitive measurement of dimerization (Chung et al., 2018, 2019; Kaizuka & Groves, 2004; Knight & Falke, 2009). If Grb2 increases the population of Btk dimers, we would expect to see a decrease in the diffusion constant of individual complexes on the membrane in the presence of Grb2. If Grb2 increases the affinity of Btk for the membrane, this would be manifested as an increase in Btk dwell time, the time that single molecules of Btk stay at the membrane, through a reduction in off-rate.

**Figure 5.**
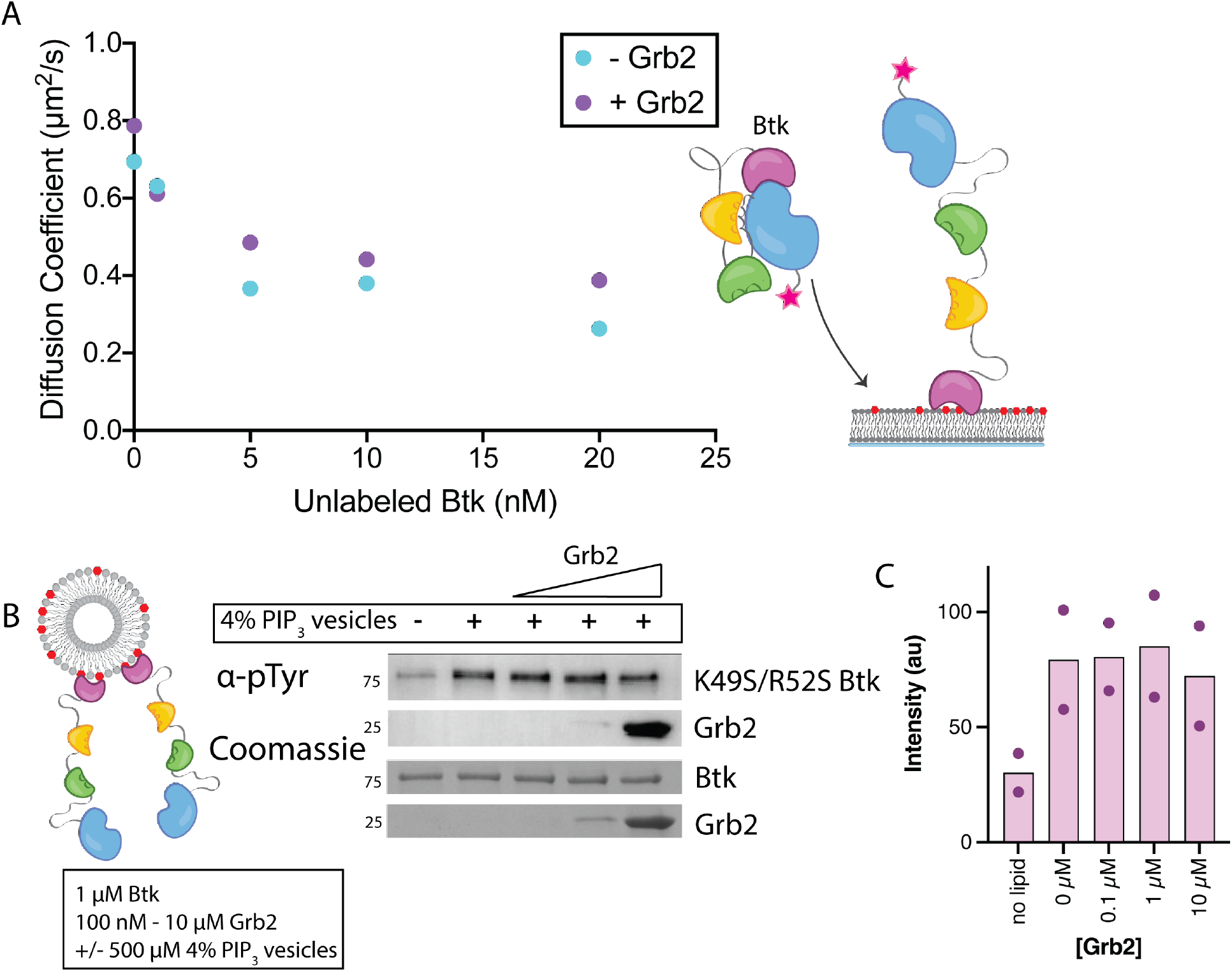
Grb2 does not enhance the dimerization of Btk on membranes. A) Diffusion constants for membrane bound Btk were determined by single-molecule tracking. Diffusion constants are plotted as a function of solution concentration of unlabeled Btk for conditions with or without Grb2. B) Western blot of Btk K49S/R52S (PH-TH peripheral PIP_3_ binding site mutant) phosphorylation with 4% PIP_3_ while titrating the amount of Grb2. See Figure 5 Supplement 2 for independent experiment replicate. C) Quantification of Btk K49S/R52S phosphorylation measured from (B), along with one independent experiment replicate, bar represents the mean intensity.

To enable site-specific labeling of full-length Btk we used unnatural amino acid incorporation of an azido-phenylalanine (AzF) group on the surface of the kinase domain of Btk (Chatterjee, Sun, Furman, Xiao, & Schultz, 2013; Chin et al., 2002). AzF enables the use of an azide-reactive dye to label the protein and eliminates non-specific labeling at other sites (see Materials and Methods for details). Several sites were tested, and incorporation of AzF at position 403 (Thr 403 in wild-type Btk) showed the best yield of labeled Btk. Thr 403 is a surface-exposed sidechain in the N-lobe of the kinase domain, and we do not anticipate that incorporation of AzF at this position will disturb the structure of Btk. For all data utilizing fluorescent full-length Btk, the construct used is Btk T403AzF labeled with azide reactive Cy5 (Btk-Cy5).

The surface density of Btk-Cy5 on the membrane was observed to increase when the solution concentration of unlabeled Btk was increased from 0 nM to 20 nM, in the presence of very low concentrations of Btk-Cy5 (500 pM-1 nM). This enables the monitoring of single molecules of Btk-Cy5, which was done either in the presence of Grb2 at 50 nM bulk concentration, or without Grb2. From each sample, step-size distributions were compiled from the single-molecule trajectories to assess the various time-dependent components to Btk diffusion on the membrane. Step-size distributions generally required fitting to three components, while two-component exponential fits were sufficient for the binding dwell time data. Both with and without Grb2, the fastest of the three diffusion constants (corresponding to monomeric Btk) decreases with increasing Btk concentration, indicating the formation of Btk dimers. The presence of Grb2 does not change the diffusive behavior of Btk at any of the concentrations used in these experiments. The decrease in Btk diffusion constant with increasing solution concentration of Btk is consistent with what is observed for the Btk PH-TH module at these concentrations, confirming that full-length Btk interacts with the membrane in the same way as does the Btk PH-TH module (Figure 5A and Figure 5 – supplement 1) (Chung et al., 2019). These experiments show that Grb2 does not change the dynamics of membrane-bound Btk, either through changes in the dimer population or changes in the membrane affinity.

We checked to see if the activity of a Btk mutant, K49S/R52S, which is incapable of significant dimerization on the membrane, could be rescued by the addition of Grb2. The K49S/R52S mutations eliminate a PIP_3_ binding site and prevent dimerization of the PH-TH module (Chung et al., 2019). A Western blot analysis shows no change in phosphorylation of the mutant Btk upon addition of Grb2, over a range of Grb2 concentrations (Figure 5B-C and Figure 5 – supplement 2).

### Grb2 can recruit Btk to clusters of scaffold proteins

We studied how the ability of Grb2 to bind to Btk might impact the localization of Btk on the membrane, by monitoring the interaction of Btk with the scaffold protein LAT on supported-lipid bilayers. LAT is similar to the B cell scaffolding protein SLP65/BLNK. Our use of LAT, rather than SLP65/BLNK, was predicated by our extensive prior work with LAT on supported-lipid bilayers (Hashimoto et al., 1999; Huang et al., 2019, 2016; Huang, Chiang, et al., 2017; Huang, Ditlev, et al., 2017; Koretzky et al., 2006; Su et al., 2016).

LAT signaling clusters can be generated from minimal components on supported-lipid bilayers. Bilayers containing 4% DGS-NTA(Ni) were prepared with both LAT and the Src-family kinase Hck tethered to the membrane through His-tags. LAT was phosphorylated by Hck, as described previously, before other components were added (Figure 6A) (Huang et al., 2019, 2016; Huang, Ditlev, et al., 2017). The diffusion constants for phosphorylated LAT and Hck are similar to those for the lipids under these conditions, indicating lack of clustering. Upon the addition of Grb2 and the proline-rich region of the Ras activator SOS (SOS-PRR), the phosphorylated LAT undergoes a protein condensation phase transition and forms gel-like domains of protein-rich areas in which the LAT no longer diffuses freely (Figure 6B) (Huang, Ditlev, et al., 2017; Su et al., 2016). This phase transition is thought to be analogous to the formation of LAT signaling clusters in T cells (Ganti et al., 2020). Using this system, we checked whether Btk could be recruited into the reconstituted LAT clusters through its interaction with Grb2.

**Figure 6.**
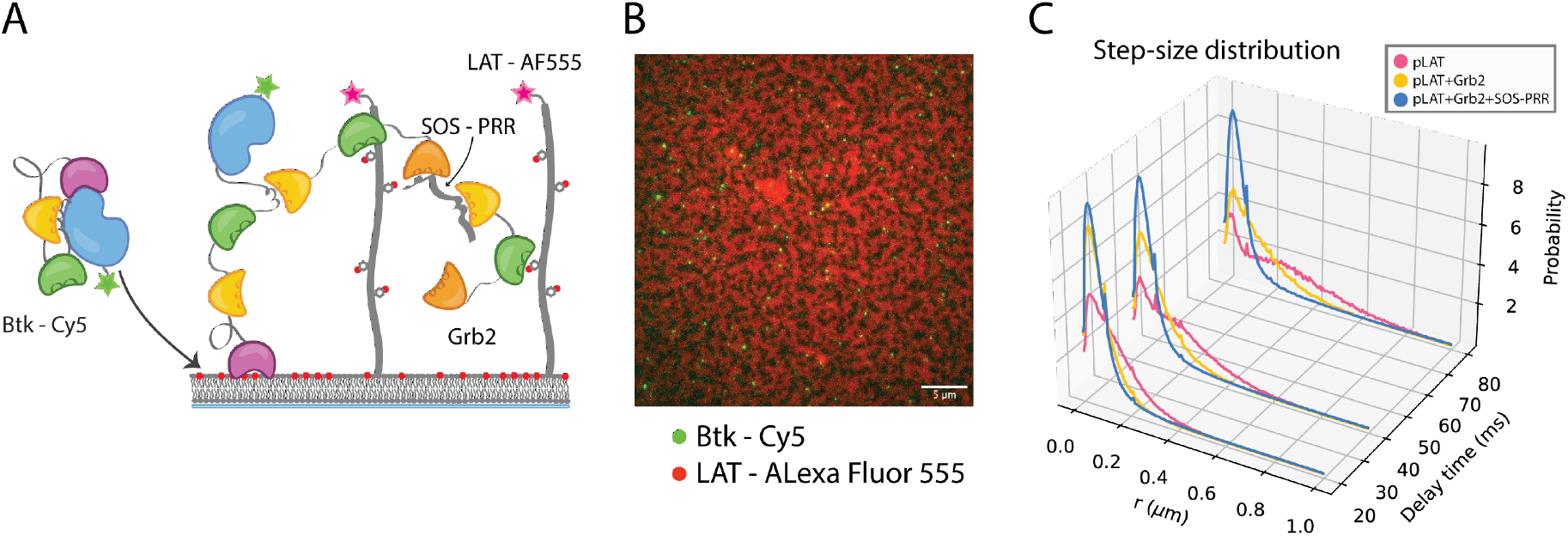
Btk can be recruited to scaffold proteins through interaction with Grb2. A) Cartoon schematic of predicted mechanism for Btk recruitment into LAT signaling clusters. B) Overlay of image of Btk-Cy5 (green) and LAT-Alexa Fluor 555 (red) after LAT phase transition. C) Step-size distribution for Btk-Cy5 under each condition: phospho-LAT, phospho-LAT + Grb2, phase transitioned LAT (phospho-LAT + Grb2 + SOS-PRR). The step-size distribution was calculated at multiple delay times. Six different positions across the bilayer were analyzed of 500-600 frames, and one independent experiment was used to confirm trends observed here.

Btk was reconstituted on supported-lipid bilayers containing 4% PIP_3_ and DGS-NTA(Ni) lipids. LAT and Hck were tethered to these bilayers as well, as described above, and LAT was phosphorylated by Hck. Subsequently, the components of the LAT signaling cluster were added sequentially along with Btk. In one condition, Btk-Cy5 was added alone to the phosphorylated LAT containing bilayers, in another condition Grb2 and Btk-Cy5 were added together, and a third condition involved by Grb2, SOS-PRR and Btk-Cy5 (Figure 6A). For the final condition, the bilayers were allowed to incubate for 1 hour to promote formation of the condensed-phase LAT domains (Figure 6B). Fluorescently labeled SOS-PRR cannot be recruited to supported-lipid bilayers containing Btk alone (Figure 6 – supplement 1).

Single molecules of Btk-Cy5 were tracked at several different positions on the bilayer for each condition. These tracks were then used to compile a step-size distribution based on all Btk-Cy5 tracks for a given condition, as described (J. J. Lin et al., 2020). The step-size distributions reflect the diffusive behavior of the entire population of Btk under a given condition; the shorter the step-sizes, the slower moving the Btk molecules. When Btk was incubated with phosphorylated LAT alone the step-size distribution is similar to that obtained for low concentrations of Btk alone on 4% PIP_3_-containing supported-lipid bilayers (Figure 6C).

Addition of Grb2 shifts the step-size distribution to shorter steps, suggesting two possible situations. One possibility is that, Grb2 is able to simultaneously bind Btk and LAT, thus slowing Btk molecules through an additional anchor point to the membrane (via the Grb2-LAT complex). The second possibility is that Grb2 alone promotes condensation of LAT (C.‑W. Lin et al., 2021), creating small domains of dense LAT, within which Btk cannot diffuse freely. The addition of SOS-PRR along with Grb2 induces the full phase transition of the phosphorylated LAT. The step-size distribution shifts even further left under this condition, suggesting that Btk has been trapped within the LAT dense phase (Figure 6C). This observation suggests that Btk is likely to be tethered to the phosphorylated LAT molecules through binding of the SH2 domain of Grb2 to phosphotyrosine residues on LAT and binding of the SH3 domains of Grb2 to Btk. This interaction leads to recruitment of Btk into the LAT condensate.

### Ideas and speculation

The Tec kinases, including Btk and Itk, constitute one of three families of cytoplasmic tyrosine kinases that contain a conserved SH3-SH2-kinase unit known as the Src module (Shah et al., 2018; Yeung et al., 2021). The other two kinase families that contain a Src module are the Src-family kinases and the Abl family. The essential regulatory mechanism operative in these kinases is an interaction between the SH3 domain and the linker connecting the SH2 domain to the kinase domain – this interaction stabilizes the kinase domain in an autoinhibited conformation, and displacement of the SH3 domain activates these kinases very strongly (Moarefi et al., 1997).

The three Src-module families differ in terms of how additional elements in the kinase support the autoinhibitory interaction between the SH3 domain and the SH2-kinase linker. The kinase activity of the Src-family kinases is suppressed by phosphorylation of the C-terminal tail, which organizes the SH3-SH2 unit into the inhibitory conformation. Abl is regulated by an N-terminal capping segment that presents a myristoyl group that helps anchor the SH2 domain to the kinase domain (Hantschel et al., 2003; Nagar et al., 2003).The N-terminal PH-TH module shared by all Tec-family kinases is a critical and distinguishing element (Andreotti, Joseph, Conley, Iwasa, & Berg, 2018; Devkota, Joseph, Boyken, Fulton, & Andreotti, 2017; Joseph et al., 2017; Wang et al., 2015). In Btk, dimerization of the PH-TH module on membranes is essential for activation, and this dimerization is promoted by the binding of the PH-TH module to PIP_3_ in the membrane (Chung et al., 2019; Wang et al., 2019, 2015). This mechanism is not a universal feature of Tec kinase regulation – the PH-TH module of Itk, for example, does not undergo dimerization on the membrane (Chung et al., 2019).

In this paper we present the discovery of an unexpected role for Grb2 in the control of Btk activity. We show that Grb2 can bind and enhance the kinase activity of Btk in the presence of PIP_3_. Previous studies have shown that the Grb2 N-terminal SH3 domain could bind Btk through interaction with the mIgG tails and SLP65 and potentiate downstream signaling (Engels et al., 2014; Kurosaki & Tsukada, 2000). We propose that Grb2 binding could activate Btk by displacing the Btk SH3 domain and releasing the inhibitory contacts within the Src module of Btk (Figure 7). Elucidating the details of the activation mechanism requires further study. We demonstrate that Btk kinase activity increases with the addition of Grb2 only with simultaneous availability of PIP_3_, which is an important point because Grb2 is a ubiquitous protein that is highly expressed in many cell lines (Shi et al., 2016). This ensures that the Btk signal remains responsive to activation of the B cell receptor.

**Figure 7.**
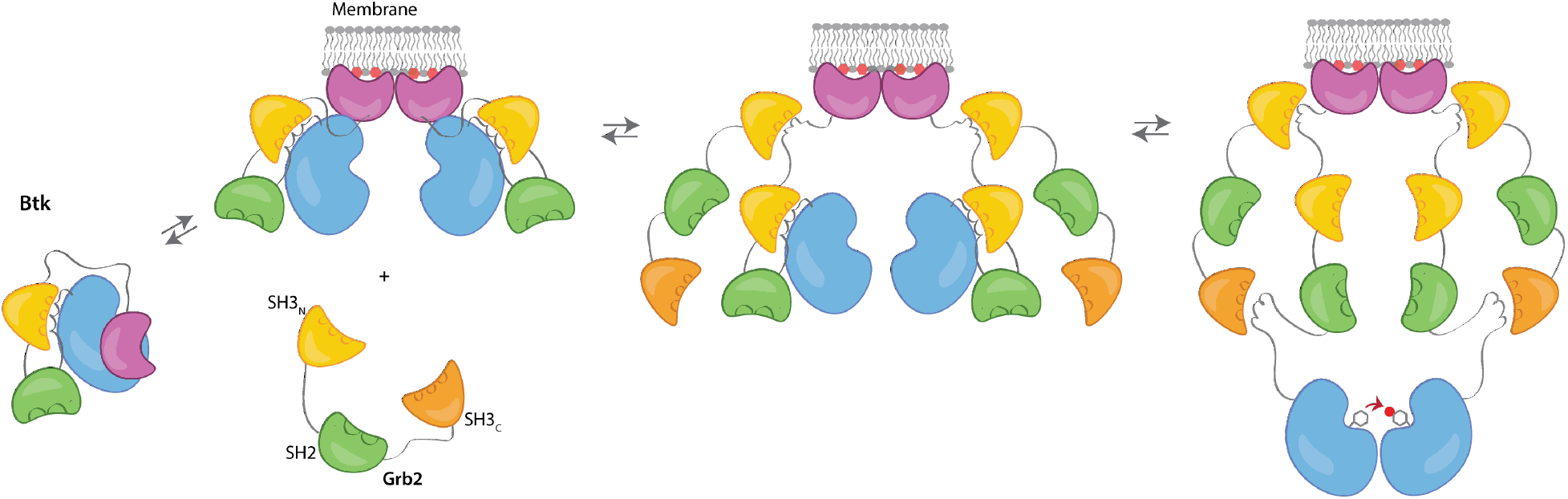
Grb2 enhances Btk activation at the membrane. Our data are consistent with a model in which PIP_3_ binding at the membrane is not sufficient for full activation, and in fact Btk is still able to maintain some autoinhibition after membrane recruitment. Upon recruitment of Grb2 to the proline-rich linker of Btk, the second Grb2 SH3 domain is able to bind the SH2-kinase linker of Btk and displace Btk’s SH3 domain, resulting in full release of autoinhibition.

A corollary of our findings is that membrane localization of Btk is insufficient for maximal stimulation of Btk activity – the interaction of Btk with Grb2 at the membrane causes increased Btk activity and stronger phosphorylation of PLCγ2. This fact may help reconcile discrepancies between NMR studies of Btk autoinhibition (Joseph et al., 2017) and inferences drawn from a crystal structure of a PH-TH-kinase construct of Btk (Wang et al., 2015). In the crystal structure, the PH-TH module is docked on the C-helix in the N-lobe of the kinase domain, and appears to stabilize an inactive conformation of the kinase domain. NMR analysis of Btk in solution shows, however, that the interaction between the PH-TH module and the kinase domain involves the activation loop, not helix C. An important point is that the crystal lattice is actually formed by dimers of Btk, with the PH-TH module forming the canonical Saraste dimer while docked on the C-helix of the inactive kinase domain (PDB 4Y93) (Wang et al., 2015). Thus the crystal structure might represent a dimeric and autoinhibited form of Btk at the membrane, with the PH-TH module forming the Saraste dimer. Grb2 might then serve to detach the PH-TH module from the kinase domain, by binding the proline-rich linker (Figure 7). Testing such a model is an important direction for future work.

By looking directly at Btk phosphorylation in the presence of Grb2 in our reconstituted system, we reveal an additional consequence of Grb2 binding – Grb2 binding can recruit Btk to signaling clusters at the membrane. In particular, the increased phosphorylation by Btk that we observe for a segment of PLCγ2 in the presence of Grb2 shows how Grb2 binding of Btk could have direct impact on the downstream signaling of Btk, subsequently increasing the population of active PLCγ2 molecules at the membrane. This is in line with observations in cells (Chiu, Dalton, Ishiai, Kurosaki, & Chan, 2002; Engels et al., 2014; Hashimoto et al., 1999; Kurosaki & Tsukada, 2000). In the absence of Grb2, Btk activation is slow, even when PIP_3_ levels are high and can promote dimerization. In the presence of Grb2, Btk phosphorylation proceeds much more rapidly. This work illuminates a new level of regulation within Btk, in which optimal signaling may rely on interaction from an adaptor molecule that both stimulates activity and facilitates localization of Btk with downstream substrates.

## Materials and Methods

### Construct definitions

| **Construct** | **Gene** | **Vector** | **N-terminal Tag** |
| --- | --- | --- | --- |
| Grb2 | Full-length, UniProt P62993 | pET28 | His<sub>6</sub>-TEV (Tobacco etch virus) site |
| Btk | Full-length, UniProt Q06187 | pET28 | His<sub>6</sub>-SUMO |
| Btk SH3-SH2-kinase | Residues 212-659 | pET28 | His<sub>6</sub>-SUMO |
| Btk SH2-Kinase | Residues 281-659 | pET28 | His<sub>6</sub>-SUMO |
| Btk kinase | Residues 402-659 | pET28 | His<sub>6</sub>-SUMO |
| Btk proline-rich linker | Residues 171-214 | pET28 | His<sub>6</sub>-SUMO |
| PLCγ2-peptide fusion | PLCγ2 Residues 746-766 UniProt P16885, EGFP | pET28 | His<sub>6</sub>-SUMO |
| Grb2 SH3<sub>N</sub>-SH2 | Residues 1-153 | pET28 | His<sub>6</sub>-TEV site |
| Grb2 SH2-SH3<sub>C</sub> | Residues 60-217 | pET28 | His<sub>6</sub>-TEV site |
| Grb2 SH3<sub>N</sub>-SH3<sub>C</sub> | Residues 1-59, 154-217 | pET28 | His<sub>6</sub>-TEV site |
| Grb2 SH3<sub>N</sub> | Residues 1-59 | pET28 | His<sub>6</sub>-TEV site |
| Grb2 SH2 | Residues 60-153 | pET28 | His<sub>6</sub>-TEV site |
| Grb2 SH3<sub>C</sub> | Residues 154-217 | pET28 | His<sub>6</sub>-TEV site |
| Grb2 R86K | Full-length, R86K | pET28 | His<sub>6</sub>-TEV site |
| Grb2 Y160E | Full-length, Y160E | pET28 | His<sub>6</sub>-TEV site |
| Btk T403AzF | Full-length, T403Amber STOP | pET28 | His<sub>6</sub>-SUMO |
| Btk K49S/R52S | Full-length, K49S/R52S | pET28 | His<sub>6</sub>-SUMO |
| LAT | Residues 30-233, Uniprot O43561 | pET28 | His<sub>6</sub>-TEV site |
| Hck | Residues 252-520, UniProt P08631 | pET28 | His<sub>6</sub>-TEV site |
| SOS-PRR | Residues 1,051-1,333, UniProt Q07889 | pET28 | His<sub>6</sub>-TEV site |
| YopH | UniProt P15273 | pCDFDuet |  |

### Protein preparation

For preparation of full-length Btk, the plasmid was transformed into BL21(DE3) *Escherichia coli* (*E. coli*) containing the YopH expression plasmid (described above) and plated on kanamycin and streptomycin containing agar plates (Seeliger et al., 2005). Transformed cells were first grown in a 200 mL Terrific broth containing 100 μg/mL kanamycin and streptomycin overnight culture at 37°C. This was then split into 6 L of Terrific broth containing 100 μg/mL kanamycin and streptomycin and grown to an optical density of 1-1.5 at 37°C. The cultures were mixed 1:1 with new media and antibiotic at 4°C and 1 mM isopropyl β-D-1-thiogalactopyranoside (IPTG) and maintained at 4°C to grow overnight. After overnight expression, the bacteria was spun down and resuspended in 15-30 mL of Ni A buffer (500 mM NaCl, 20 mM Tris-HCl pH 8.5, 20 mM Imidazole, 5% Glycerol). These were then flash frozen and kept at −80°C until the next step of the purification.

Cell pellets were thawed and then lysed by homogenization or sonication with addition of phenylmethylsulfonyl fluoride. The lysate was then spun down at 16500 rpm for one hour. Supernatant was collected and flowed over a HisTrap FF (GE) column. The column was then washed with 10 column volumes (CV) Ni A buffer, followed by four CV Ni B buffer (500 mM NaCl, 20 mM Tris-HCl pH 8.5, 500 mM Imidazole, 5% Glycerol). The elution fraction was then loaded onto a desalting column equilibrated in Buffer A (150 mM NaCl, 20 mM Tris-HCl pH 8.0, 5% Glycerol). The protein peak was collected and incubated with ULP1 protease overnight at 4°C.

After His tag cleavage, the sample was run over a second HisTrap column, this time the flow through and wash were collected. In some cases, a gradient of immidazole was applied as the Btk PH domain has some affinity for the Ni column even in the absence of the His tag. The protein was then concentrated in an Amicon Ultra centrifugal filter (Millipore Sigma) to less than 2 mL total volume. This was then loaded onto an S200 30/300 (GE) column equilibrated in Buffer A for biochemistry or Buffer B (150 mM NaCl, 50 mM Hepes pH 7.4, 5% Glycerol) for imaging studies. Fractions containing the purest target protein were collected and concentrated. These were then aliquoted and stored at −80°C after flash freezing.

All other constructs were purified in a similar manner with the following changes. All constructs that did not contain a kinase domain were transformed into BL21(DE3) *E.* coli without the YopH plasmid, and therefore they were grown with only kanamycin. Expression for these constructs was carried out overnight at 18°C. Grb2 constructs and PLCγ2-peptide fusion express at much higher levels than the Btk constructs and therefore only 1-2 liters were prepared at a given time, and therefore no overnight culture was required. For all constructs that contain a His tag after purification (LAT, Hck, and His-tagged Btk constructs), the protease and second HisTrap column were eliminated. Grb2 constructs and SOS-PRR each contain a TEV site and therefore TEV protease was used to remove the His tag.

### Protein fluorescent labeling by maleimide conjugation

Grb2 and SOS-PRR were each prepared at a concentration of 50 μM and incubated with 5 mM DTT for 30 minutes on ice to ensure all accessible cysteines were reduced. Maleimide conjugated dye was dissolved in anhydrous DMSO and added in equimolar amounts to Grb2 or 3-fold molar excess to SOS-PRR and incubated at room temperature for 30 minutes or at 4°C overnight, depending on the stability of the protein. The reaction was then quenched with 10 mM DTT for 30 minutes at 4°C. After quenching, the protein was diluted in 10 – 15 mL fresh buffer, and then concentrated in an Amicon Ultra centrifugal filter, allowing free dye to be removed in the flow through. This process was continued until no more free dye could be easily detected in the flow through. At this point the protein was then purified by gel filtration in Buffer B as described above. Labeling efficiency was calculated based on the absorbances at 280 nm and the peak excitation wavelength for the dye, taking into account dye contribution to the 280 nm signal. A note about Grb2 labeling: the protein tends to aggregate when exposed to an excess of dye, and therefore we favored under labeling by only providing equimolar amount of dye.

### Protein fluorescent labeling by azido phenylalanine incorporation

For site specific labeling of full-length Btk for single molecule studies, we used unnatural amino acid incorporation of an Azido phenylalanine (AzF) residue (Amiram et al., 2015; Bard & Martin, 2018; Chatterjee et al., 2013; Chin et al., 2002). An amber codon (UAG) was introduced at the desired labeling position in the plasmid for Btk expression, and it was ensured that the stop codon for this gene was not amber. This plasmid was co-transformed into BL21(DE3) containing a plasmid expressing YopH with the pUltra-pAzFRS plasmid. A 5 mL starter culture was grown overnight in TB at 37°C from this transformation. This was used to inoculate 1 L TB the following morning which was grown to an optical density between 0.5 and 1 at 37°C. This culture was spun down and resuspended in 167 mL cold (4°C) TB containing 2 mM AzF, before induction, this culture was allowed to grow for 1.25 hours at 4°C. The culture was then induced with 1 mM IPTG and grown overnight at 4°C. After this, protein purification proceeded as normal until just before the gel filtration step, except that all reducing agents were left out of the buffers. Yield of the protein is drastically reduced, however for our applications a low yield was not a problem as long as labeling was feasible.

For labeling, the purified protein was concentrated to about 300 μL. 5 mM 5,5-dithio-bis-(2-nitrobenzoic acid) (DTNB, Ellman’s reagent) was prepared in 50 mM HEPES, pH 7.0, 250 mM KCl, 5% glycerol. DTNB was added at 15X molar excess to the protein solution and incubated at room temperature for 10 minutes. The protein was then cooled back to 4°C and 300 μM dibenzocyclooctyl (DBCO)-Cy5 was added from a 30 mM stock in DMSO. This was incubated overnight at 4°C. The reaction was quenched with 5 mM DTT and incubated at 4°C for 1 hour. At this point the labeled protein was purified by gel filtration, then concentrated to an appropriate concentration, flash frozen and stored at −80°C. This protocol was adapted from the lab of Andreas Martin (Bard, Bashore, Dong, & Martin, 2019; Bard & Martin, 2018; Lander et al., 2012).

### Western blot assays for Btk kinase activity

Samples were prepared with 2X concentration of each component depending on the condition to be tested (2 μM Btk, 1 mM 4% PIP_3_ single unilamellar vesicles (SUVs), 0-20 μM Grb2) in Buffer A, final concentrations during the reaction are indicated for each blot (Figure 3A, 3C, 4A, 4C and 5B). These components were incubated for 15 minutes at room temperature and then diluted 1:1 in activation buffer (20 mM MgCl_2_, 2 mM ATP, 2 mM sodium vanadate). These were mixed and the reaction proceeded at room temperature for the designated amount of time, indicated for each blot separately. The reaction was then quenched by mixing 1:1 with Quench buffer (166 mM Tris-HCl pH 6.8, 10% SDS, 10 mM DTT, 3 μM bromophenol blue, 10% glycerol, 100 mM EDTA). These samples were then immediately heated at 90°C for 15 minutes, followed by loading onto two 12 or 15% SDS page gels, which were run at 250V for 35 minutes. One gel was stained with Coomassie blue, the second was prepared for a Western blot transfer.

Filters were soaked in Western blot transfer buffer (25 mM Tris-HCl pH 7.4, 192 mM glycine) supplemented with 0.1% SDS. The membrane was activated in MeOH for 1 minute and then transferred to Western blot transfer buffer supplemented with 20% MeOH. The protein was transferred in a semi-dry apparatus to the membrane at 25 V for one hour. This membrane was then blocked in 5% Carnation non-fat milk powder for one hour while shaking gently. To probe for phospotyrosine, the membrane was then transferred to a 1:2000 dilution of α-phosphotyrosine (Cell Signaling) for shaking overnight at 4°C. The following day, the membrane was washed four times for 10 minutes each in TBS-T (20 mM Tris pH 7.5, 150 mM NaCl, 0.1% Tween-20). The membrane was then transferred to a secondary antibody solution containing a 1:5000 dilution of α-mouse HRP (Cell Signaling) for one hour of shaking at room temperature. The membrane was then washed again four times for 10 minutes each in TBS-T. At this point the blot was imaged with Western Bright (Advansta). Blots were quantified using Fiji (ImageJ). For all bands, a local background intensity was measured, and band intensity was calculated as a change in intensity as compared to its local background. For comparison of intensities across replicate blots, average background intensity was calculated for each replicate (based on the local background intensities mentioned above). A scale factor was generated by dividing background intensity of replicate 1 by the average background intensity of replicate 2. The change in intensity for replicate 1 was then divided by the scale factor to ensure that the overall intensity of each blot was consistent. These values are then plotted as “Intensity” for each graph (Figure 3D, 4B, 4D, 5C). It should be noted that this technique is only semi-quantitative as the intensity may not be linear with protein concentration, these graphs are provided as a tool to compare trends rather than exact values.

### Supported-lipid bilayer preparation

Vesicles were prepared by first mixing the desired ratios of PIP_3_ (Echelon Biosciences, Inc.), 1,2-dioleoyl-sn-glycero-3-[(N-(5-amino-1-carboxypentyl)iminodiacetic acid)succinyl] (nickel salt) (DGS-NTA(Ni)), and 18:1 1,2-dioleoyl-sin-glycero-3-phosphocholine (DOPC) (Avanti Polar Lipids). For experiments with full-length Btk (no His tag) 4% PIP_3_, 96% DOPC mixtures were prepared. For experiments using His-tagged Btk constructs 4% DGS-NTA(Ni), 96% DOPC mixtures were prepared. And for experiments involving LAT 4% PIP_3_, 4% DGS-NTA(Ni), 92% DOPC bilayers mixtures were prepared. All lipids were stored in solution with chloroform, except PIP_3_ which was stored in powder form and dissolved in a solution of 1:2:0.8 Chloroform:Methanol:Water just before use. After mixing, the solution was dried in an etched 50 mL round bottomed flask by rotary evaporation for 15 minutes and then under nitrogen for 15 minutes. Dried lipids could then be kept overnight at 4°C, sealed from air or used immediately.

The lipids were rehydrated in water to 1 mg/mL lipid concentration by vortexing. Single unilamellar vesicles (SUVs) were created by sonication (Analis Ultrasonic Processor 750 watt, 20 kHz) of the lipids at 33% power, 20 seconds on, 50 seconds off for 1 minute 40 seconds of active sonication. A solution of 0.25 mg/mL SUVs in 0.5X TBS was prepared. These were added to a Sticky Slide VI 0.4 (Ibidi GMBH) attached to a Piranha-etched glass slide, 80 μL of SUVs per well. The SUVs were incubated at room temperature for 30 minutes. Each well was then washed with 1 mL of Hepes Buffered Saline (HBS) under vacuum, being careful to avoid introduction of air bubbles to the chamber. When using DGS-NTA(Ni), the wells were then incubated with 100 mM NiCl_2_ for 5 minutes, and again washed with 1 mL HBS. At this point, the wells were blocked with either 1 μg/mL Poly(L-Lysine)-PEG (PLL-PEG) for 1 minute, or 1 mg/mL Bovine Serum Albumin (BSA) or β-Casein for 10 minutes. The blocking agent for each experiment was optimized by checking the mobile fraction of surface bound particles. In the case of full-length Btk on 4% PIP_3_ bilayers, 1 μg/mL PLL-PEG yielded the best results. For LAT containing bilayers, 1 mg/mL BSA was used to block. For all other experiments, 1 mg/mL β-Casein yielded the best results.

For experiments where DGS-NTA(Ni) lipids were used to coordinate His-tagged protein to the bilayers, the protein was added at the desired incubation concentration and left at room temperature for 40 minutes. The chambers were then washed with 600 μL HBS gently by hand and incubated for 20 more minutes at room temperature. They were then again washed with 600 μL HBS. Finally, they were washed with 100 μL imaging buffer containing 100 μg/mL BSA or β-Casein (depending on the blocking agent that was used initially) and 10 mM BME in HBS.

### Monitoring membrane recruitment by fluorescence microscopy

TIRF adsorption experiments were carried out and collected on a Nikon Eclipse Ti-inverted microscope and Andor iXon camera, as described (Bhattacharyya et al., 2020). TIRF data were acquired at 15 second intervals, and data were generally displayed as a difference intensity, where the baseline for a given sample was calculated from the average of the four frames preceding Grb2 addition. Single-molecule traces were recorded as described previously (J. J. Lin et al., 2020). Btk was allowed to equilibrate with the supported-lipid bilayers for 30 minutes before imaging. Movies were recorded at intervals of 20 ms, and five or more traces of 500-600 frames were collected at various places across the sample. Fluorescent molecules in these movies were tracked using the TrackMate plugin from Fiji (Image J). Tracking parameters were kept consistent across experiments. An immobile fraction of fluorescent Btk was always observed, and this fraction was excluded from the analysis. Tracks were analyzed by calculating a step size distribution and dwell time for the particles, as described in the main text.

### Reconstitution of LAT phase preparation on supported-lipid bilayers

LAT was reconstituted on supported-lipid bilayers as described (Huang, Ditlev, et al., 2017). Supported-lipid bilayers were prepared as described above with 30 nM His_6_-Hck and 150 nM His_6_-LAT-Alexa Fluor 555 on a 4% DGS-NTA(Ni), 96% DOPC. The LAT was phosphorylated by including 1 mM ATP and 10 mM MgCl_2_ in the imaging buffer and incubating for 20 minutes before adding other components. After phosphorylation, the LAT phase transition was induced by adding 6 μM Grb2 and 1.8 μM proline-rich region of SOS (SOS-PRR) and incubated for at least 1 hour. For experiments monitoring Btk diffusion and dwell time, 1 nM Btk T403AzF-Cy5 was also added at this point. After LAT was observed to undergo phase separation, multiple traces of Btk diffusion were recorded and analyzed as described above for single molecule tracking.

## Acknowledgements

We thank Jean Chung, William Huang, and Josh Cofsky for their help in providing protocols and troubleshooting assistance. We thank Darren McAffee and Kiera Wilhelm for sharing their data analysis pipelines for the single molecule studies, and Chun-Wei Lin, Joey DeGrandchamp, and Nugent Lew for sharing protein that was used for some experiments in this paper. We are grateful to Neel Shah, Jeanine Amacher, and Helen Hobbs for helpful conversations about kinase signaling. Finally, we thank Susan Marqusee and David Wemmer for their feedback on this project.

---

**Figure 3 Supplement 1.**
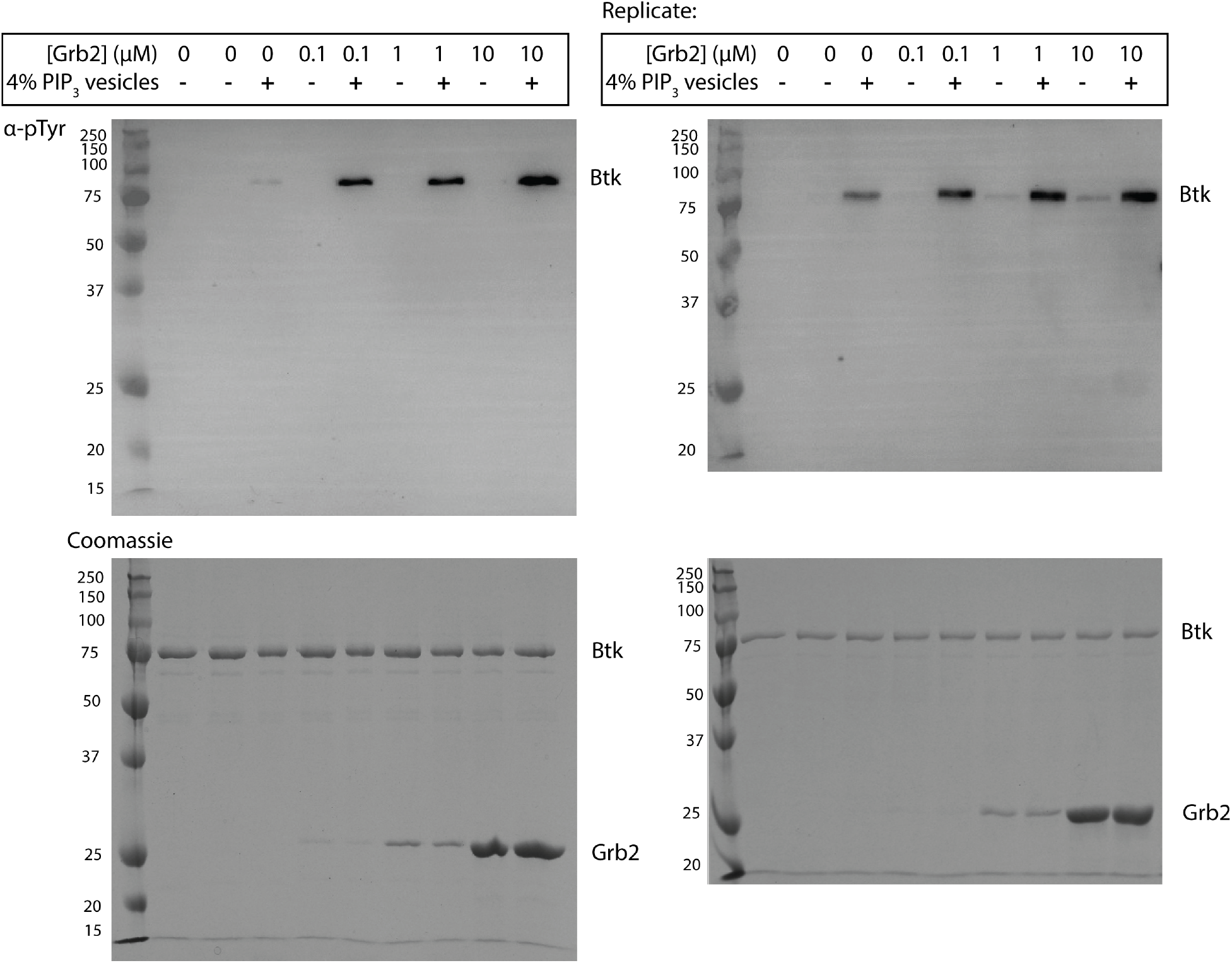
Grb2 enhances *trans-* autophosphorylation of Btk in the presence of PIP_3_ containing vesicles. Entire blot from Figure 3A in replicate, looking at autophosphorylation of Btk. For each lane 1 μM Btk was activated for 5 min in the presence or absence of 500 μM lipids, single unilamellar vesicles containing 4% PIP_3_, and the indicated concentrations of Grb2. Each experiment was carried out in at least duplicate.

**Figure 3 Supplement 2.**
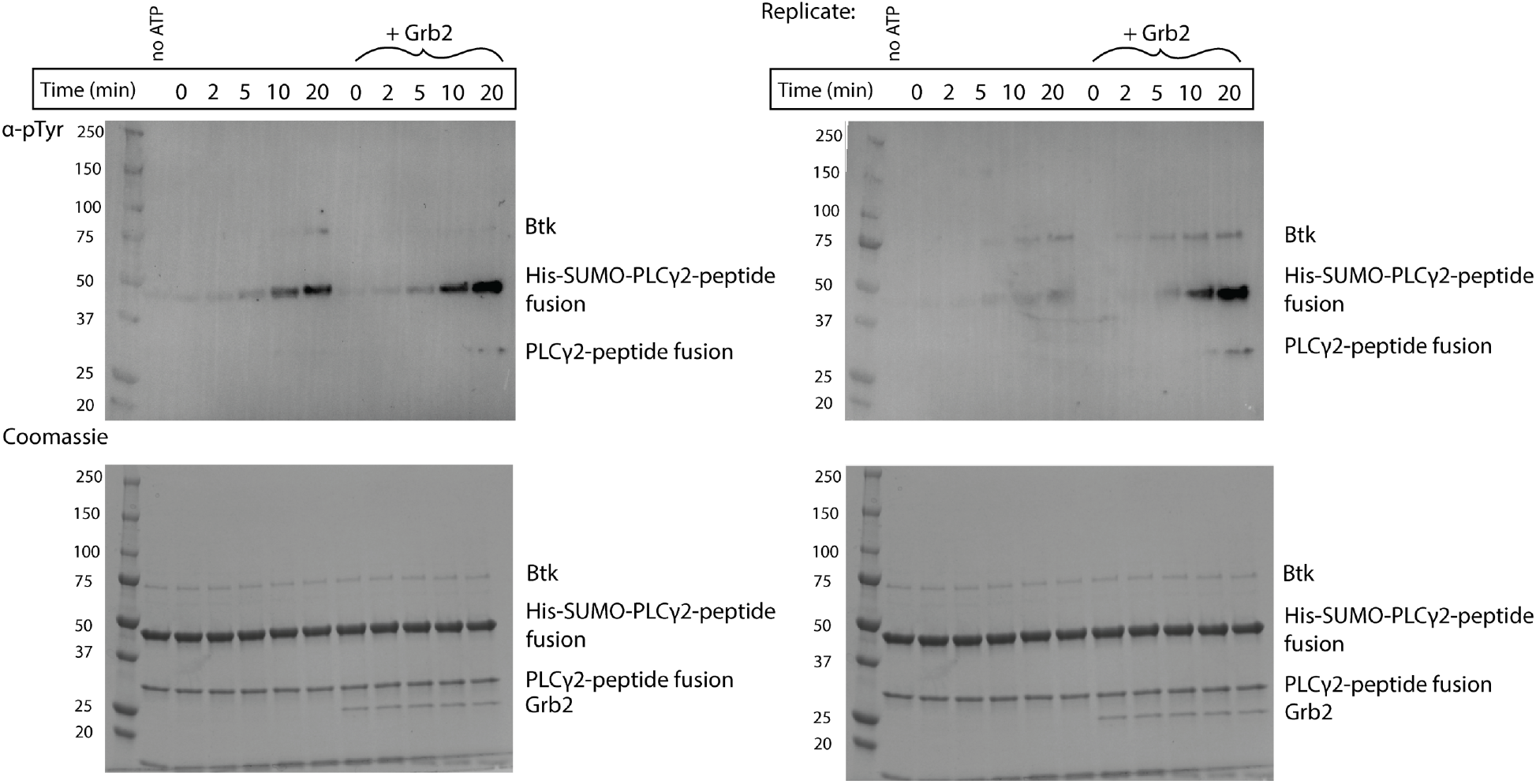
Grb2 enhances phosphorylation of PLCγ2 by Btk in the presence of PIP_3_ containing vesicles. Entire blot from Figure 3C in replicate, looking at phosphorylation of PLCγ2-peptide fusion. 1 μM Btk was activated for the indicated amount of time in the presence of 500 μM lipids, single unilamellar vesicles containing 4% PIP_3_, 10 μM PLCγ2 peptide fusion with or without 1μM Grb2. Each experiment was carried out in at least duplicate.

**Figure 4 Supplement 1.**
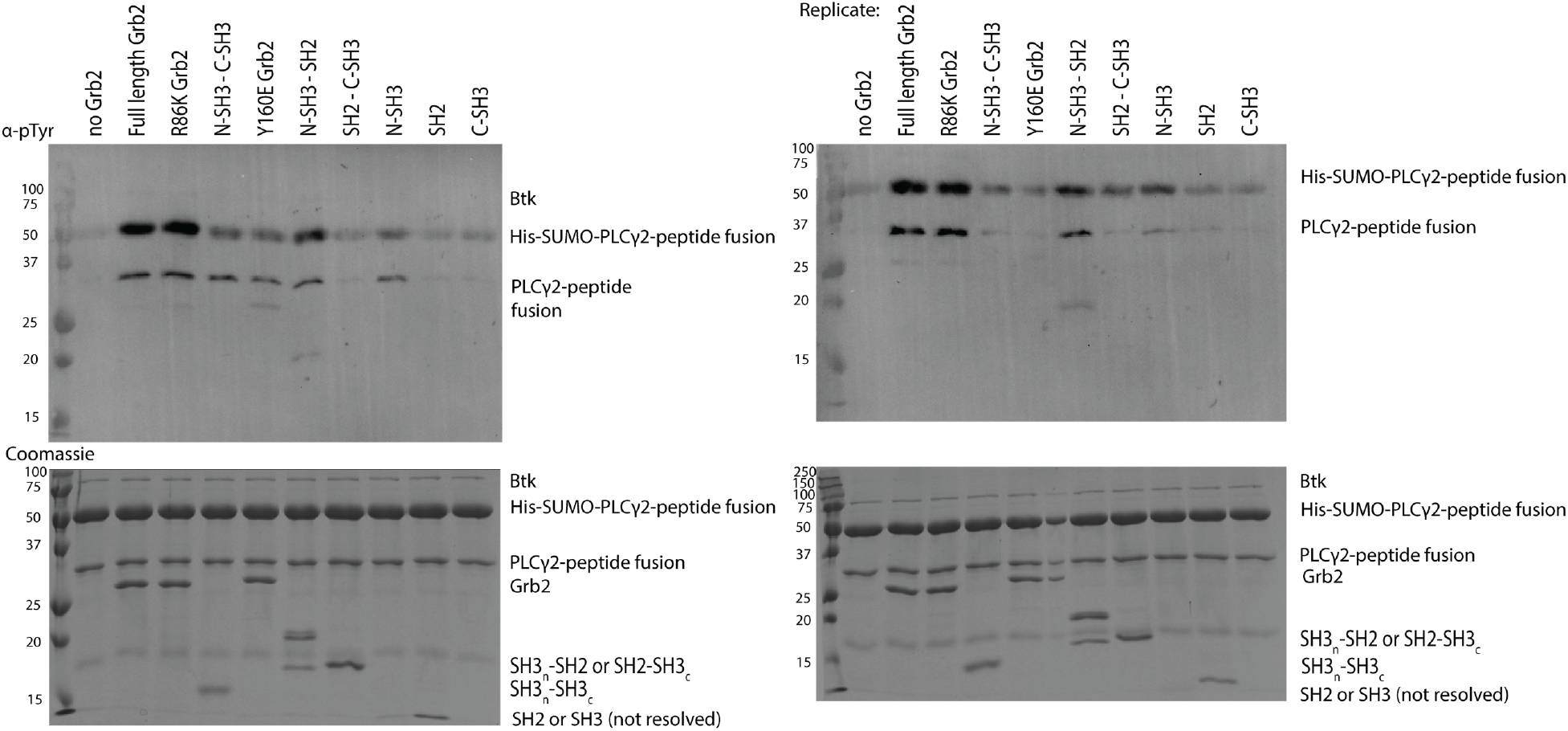
Phosphorylation of PLCγ2 by Btk with the addition of various Grb2 constructs. Entire blot from Figure 4A and replicate, looking at phosphorylation of PLCγ2-peptide fusion in the presence of Btk and varying constructs of Grb2. 1 μM Btk was activated for 10 minutes in the presence of 500 μM lipids, single unilamellar vesicles containing 4% PIP_3_, 10 μM PLCγ2 peptide fusion with 10 μM of the indicated construct of Grb2. Each experiment was carried out in at least duplicate.

**Figure 4 Supplement 2.**
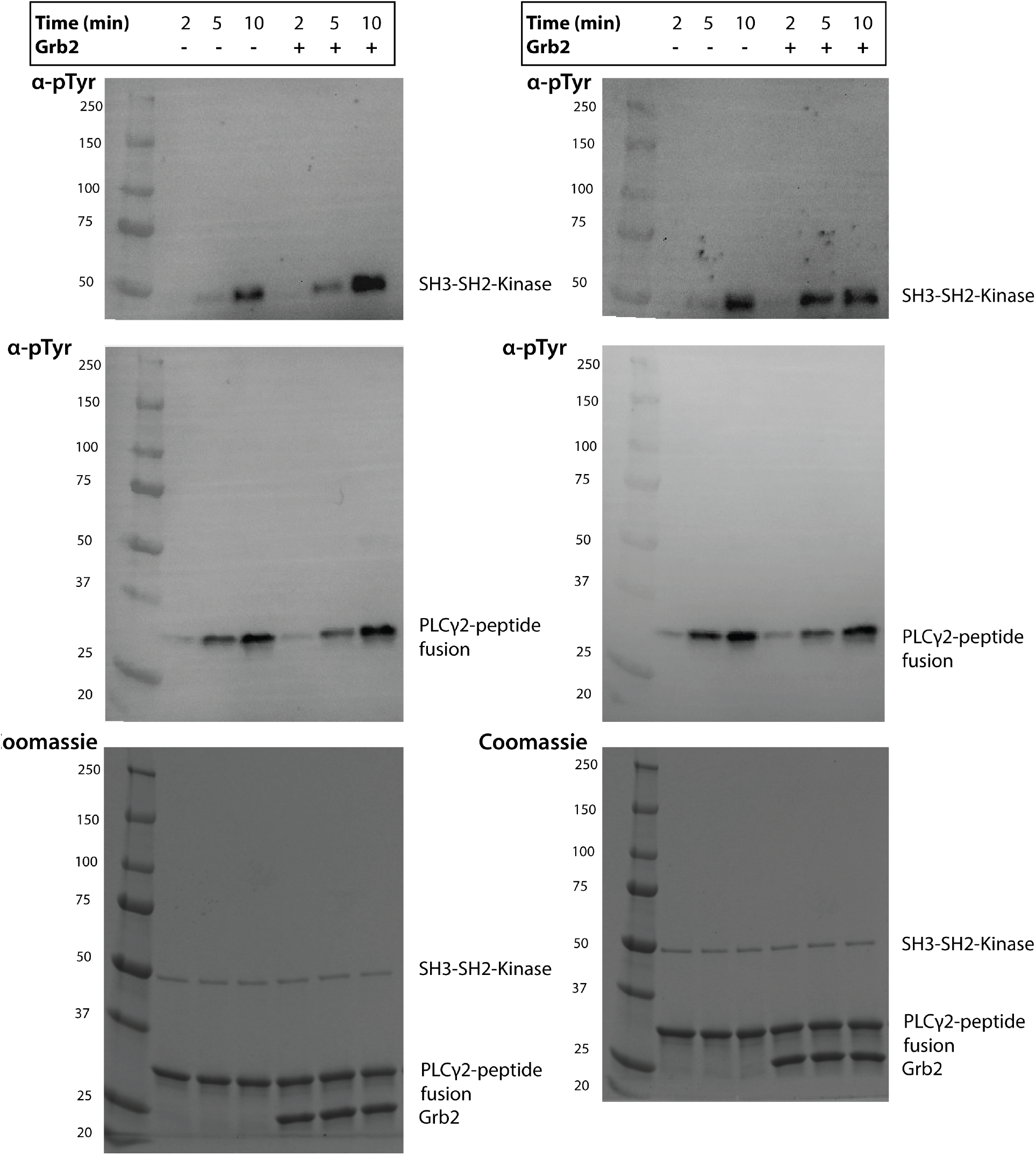
The Src module of Btk shows moderate enhancement of catalytic activity upon addition of Grb2. Entire blot from Figure 4C and replicate, looking at phosphorylation of the Src-module of Btk with Grb2 and PLCγ2-peptide fusion. 1 μM Src-module of Btk was activated for the indicated amount of time in the presence of 10 μM PLCγ2 peptide fusion with 10 μM of Grb2. Each experiment was carried out in at least duplicate.

**Figure 5 Supplement 1.**
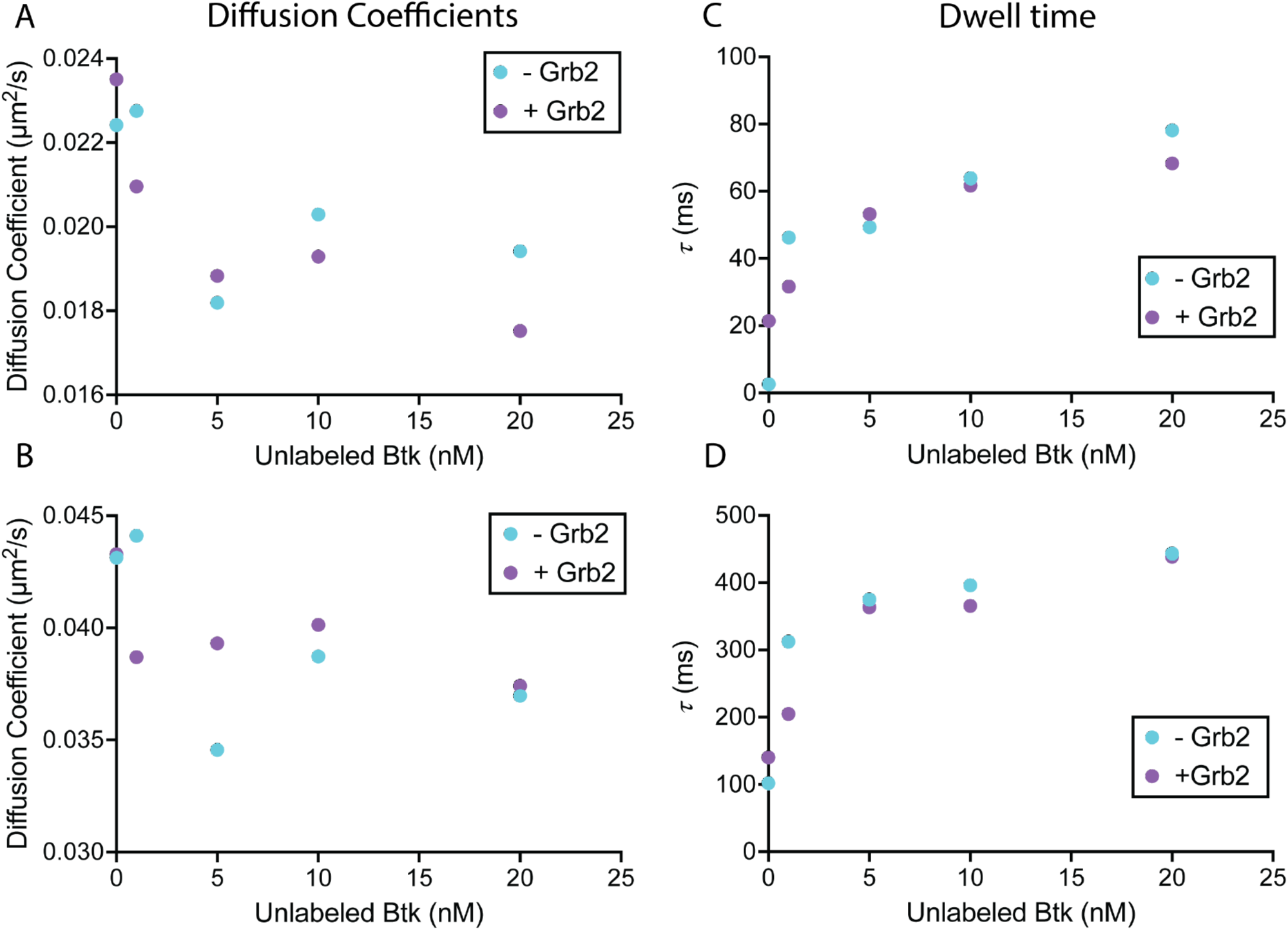
The diffusion of Btk on membranes in the absence and presence of Grb2. Remaining components of Btk diffusion as fit by single molecule tracking. A) The slowest component of the diffusion, trending slightly with increasing surface density of Btk. B) The middle component of the diffusion, remaining relatively constant across Btk concentrations. C) And D) are both components of the dwell time, which both show dependence on Btk surface density.

**Figure 5 Supplement 2.**
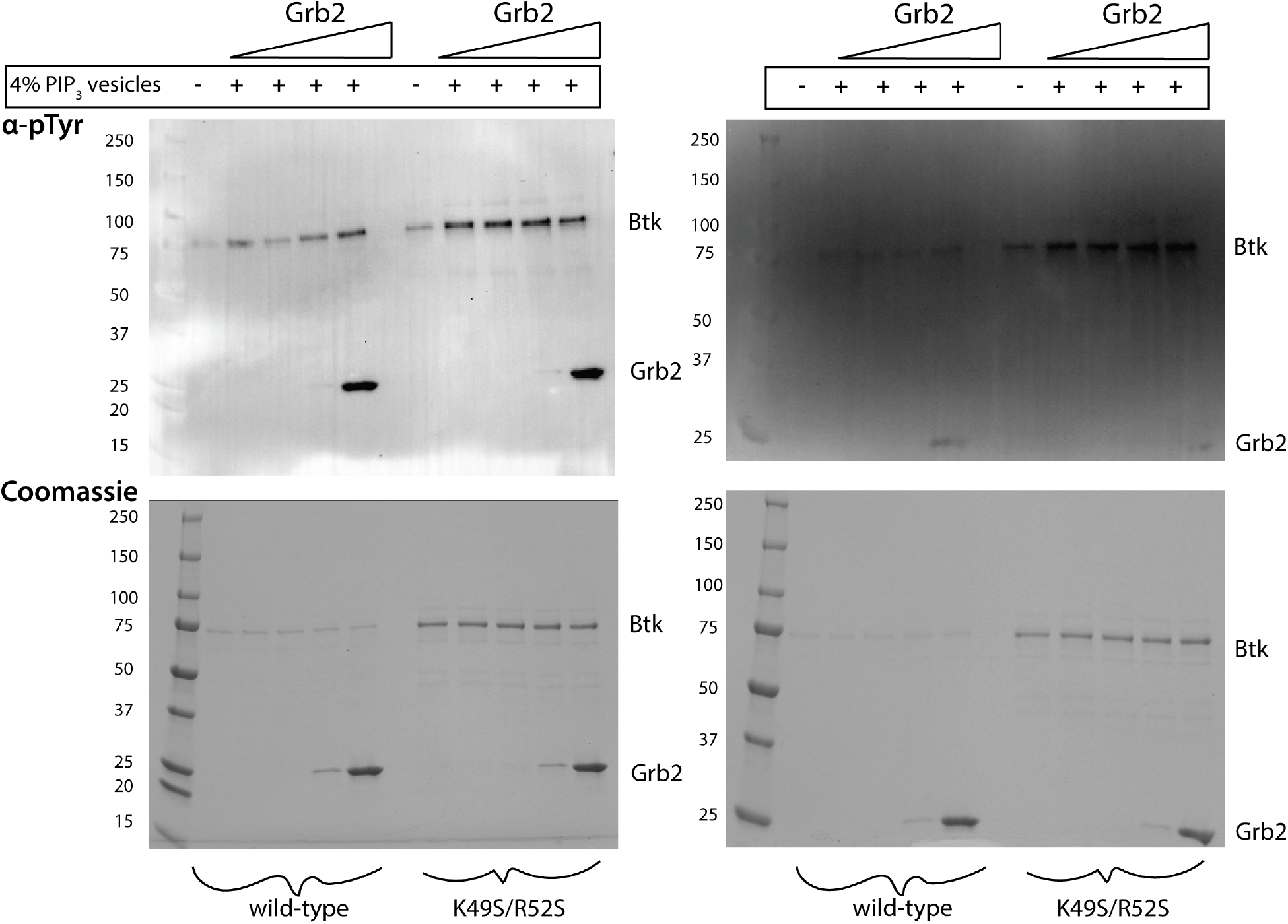
Phosphorylation of Btk K49S/R52S is not enhanced by Grb2. Entire blot from Figure 5B and replicate looking at phosphorylation of Btk K49S/R52S with the addition of Grb2. 1 μM Btk wild-type or K49S/R52S was activated for 10 minutes in the presence or absence of 500 μM lipids, single unilamellar vesicles containing 4% PIP_3_, and 100 nM - 10 μM of Grb2, each consecutive lane representing an increase in the Grb2 concentration by one order of magnitude. Each experiment was carried out in at least duplicate.

**Figure 6 Supplement 1.**
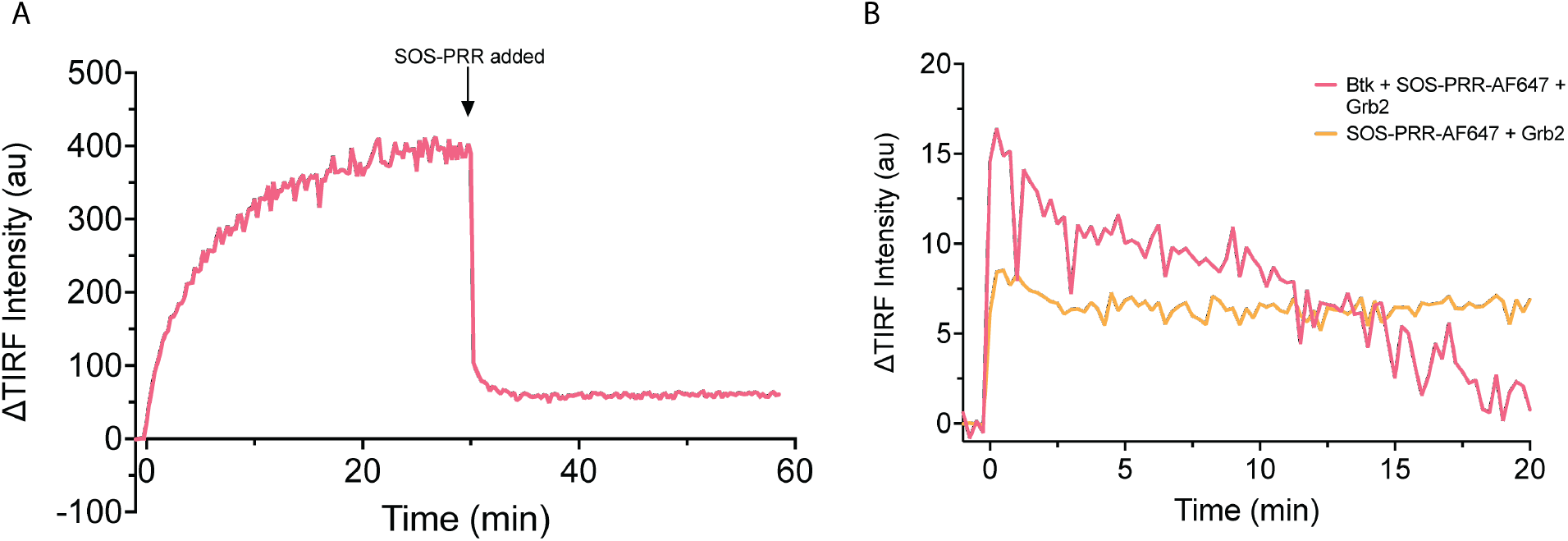
Behavior of Btk on PIP3-containing membranes with the addition of SOS proline rich region. A) Here Grb2-AlexaFluor647 and Btk have been added to a PIP_3_-containing membrane at time 0, AlexaFluor647 fluorescence intensity is measured. After 30 minutes SOS-PRR is added at the same concentration that is used in Figure 6. SOS-PRR appears to compete with Btk for Grb2 binding. B) Btk, Grb2, and SOS-PRR-AlexaFluor647 or Grb2 and SOS-PRR-AlexaFluor647 are added to PIP_3_ containing membrane and AlexaFluor647 fluorescence intensity is measured. SOS-PRR does not interact with Btk on supported-lipid bilayers.

## References

Ahmed, Z., Timsah, Z., Suen, K. M., Cook, N. P., Lee, G. R., Lin, C.-C., … Ladbury, J. E. (2015). Grb2 monomer-dimer equilibrium determines normal versus oncogenic function. Nature Communications, 6, 7354. doi: 10.1038/ncomms8354l

Amatya, N., Lin, D. Y.-W., & Andreotti, A. H. (2019). Dynamic regulatory features of the protein tyrosine kinases. Biochemical Society Transactions, 47(4), 1101–1116. doi: 10.1042/BST20180590

Amiram, M., Haimovich, A. D., Fan, C., Wang, Y.-S., Aerni, H.-R., Ntai, I., … Isaacs, F. J. (2015). Evolution of translation machinery in recoded bacteria enables multi-site incorporation of nonstandard amino acids. Nature Biotechnology, 33(12), 1272–1279. doi: 10.1038/nbt.3372

Andreotti, A. H., Joseph, R. E., Conley, J. M., Iwasa, J., & Berg, L. J. (2018). Multidomain control over TEC kinase activation state tunes the T cell response. Annual Review of Immunology, 36, 549–578. doi: 10.1146/annurev-immunol-042617-053344

Baraldi, E., Djinovic Carugo, K., Hyvönen, M., Surdo, P. L., Riley, A. M., Potter, B. V., … Saraste, M. (1999). Structure of the PH domain from Bruton’s tyrosine kinase in complex with inositol 1,3,4,5-tetrakisphosphate. Structure, 7(4), 449–460. doi: 10.1016/s0969-2126(99)80057-4

Bard, J. A. M., Bashore, C., Dong, K. C., & Martin, A. (2019). The 26S proteasome utilizes a kinetic gateway to prioritize substrate degradation. Cell, 177(2), 286–298.e15. doi: 10.1016/j.cell.2019.02.031

Bard, J. A. M., & Martin, A. (2018). Recombinant Expression, Unnatural Amino Acid Incorporation, and Site-Specific Labeling of 26S Proteasomal Subcomplexes. Methods in Molecular Biology, 1844, 219–236. doi: 10.1007/978-1-4939-8706-1_15

Bhattacharyya, M., Lee, Y. K., Muratcioglu, S., Qiu, B., Nyayapati, P., Schulman, H., … Kuriyan, J. (2020). Flexible linkers in CaMKII control the balance between activating and inhibitory autophosphorylation. ELife, 9. doi: 10.7554/eLife.53670

Cantor, A. J., Shah, N. H., & Kuriyan, J. (2018). Deep mutational analysis reveals functional trade-offs in the sequences of EGFR autophosphorylation sites. Proceedings of the National Academy of Sciences of the United States of America, 115(31), E7303–E7312. doi: 10.1073/pnas.1803598115

Chatterjee, A., Sun, S. B., Furman, J. L., Xiao, H., & Schultz, P. G. (2013). A versatile platform for single- and multiple-unnatural amino acid mutagenesis in Escherichia coli. Biochemistry, 52(10), 1828–1837. doi: 10.1021/bi4000244

Chin, J. W., Santoro, S. W., Martin, A. B., King, D. S., Wang, L., & Schultz, P. G. (2002). Addition of p-azido-L-phenylalanine to the genetic code of Escherichia coli. Journal of the American Chemical Society, 124(31), 9026–9027. doi: 10.1021/ja027007w

Chiu, C. W., Dalton, M., Ishiai, M., Kurosaki, T., & Chan, A. C. (2002). BLNK: molecular scaffolding through ’cis’-mediated organization of signaling proteins. The EMBO Journal, 21(23), 6461–6472. doi: 10.1093/emboj/cdf658

Chung, J. K., Lee, Y. K., Denson, J.-P., Gillette, W. K., Alvarez, S., Stephen, A. G., & Groves, J. T. (2018). K-Ras4B Remains Monomeric on Membranes over a Wide Range of Surface Densities and Lipid Compositions. Biophysical Journal, 114(1), 137–145. doi: 10.1016/j.bpj.2017.10.042

Chung, J. K., Nocka, L. M., Decker, A., Wang, Q., Kadlecek, T. A., Weiss, A., … Groves, J. T. (2019). Switch-like activation of Bruton’s tyrosine kinase by membrane-mediated dimerization. Proceedings of the National Academy of Sciences of the United States of America, 116(22), 10798–10803. doi: 10.1073/pnas.1819309116

Clark, S. G., Stern, M. J., & Horvitz, H. R. (1992). C. elegans cell-signalling gene sem-5 encodes a protein with SH2 and SH3 domains. Nature, 356(6367), 340–344. doi: 10.1038/356340a0

Courtney, A. H., Lo, W.-L., & Weiss, A. (2018). TCR signaling: mechanisms of initiation and propagation. Trends in Biochemical Sciences, 43(2), 108–123. doi: 10.1016/j.tibs.2017.11.008

Devkota, S., Joseph, R. E., Boyken, S. E., Fulton, D. B., & Andreotti, A. H. (2017). An Autoinhibitory Role for the Pleckstrin Homology Domain of Interleukin-2-Inducible Tyrosine Kinase and Its Interplay with Canonical Phospholipid Recognition. Biochemistry, 56(23), 2938–2949. doi: 10.1021/acs.biochem.6b01182

Duarte, D. P., Lamontanara, A. J., La Sala, G., Jeong, S., Sohn, Y.-K., Panjkovich, A., … Hantschel, O. (2020). Btk SH2-kinase interface is critical for allosteric kinase activation and its targeting inhibits B-cell neoplasms. Nature Communications, 11(1), 2319. doi: 10.1038/s41467-020-16128-5

Engels, N., Engelke, M., & Wienands, J. (2008). Conformational Plasticity and Navigation of Signaling Proteins in Antigen-Activated B Lymphocytes. In Advances in Immunology: Vol. 97 (pp. 251–281). Elsevier. doi: 10.1016/S0065-2776(08)00005-9

Engels, N., König, L. M., Heemann, C., Lutz, J., Tsubata, T., Griep, S., … Wienands, J. (2009). Recruitment of the cytoplasmic adaptor Grb2 to surface IgG and IgE provides antigen receptor-intrinsic costimulation to class-switched B cells. Nature Immunology, 10(9), 1018–1025. doi: 10.1038/ni.1764

Engels, N., König, L. M., Schulze, W., Radtke, D., Vanshylla, K., Lutz, J., … Wienands, J. (2014). The immunoglobulin tail tyrosine motif upgrades memory-type BCRs by incorporating a Grb2-Btk signalling module. Nature Communications, 5, 5456. doi: 10.1038/ncomms6456

Gale, N. W., Kaplan, S., Lowenstein, E. J., Schlessinger, J., & Bar-Sagi, D. (1993). Grb2 mediates the EGF-dependent activation of guanine nucleotide exchange on Ras. Nature, 363(6424), 88–92. doi: 10.1038/363088a0

Ganti, R. S., Lo, W.-L., McAffee, D. B., Groves, J. T., Weiss, A., & Chakraborty, A. K. (2020). How the T cell signaling network processes information to discriminate between self and agonist ligands. Proceedings of the National Academy of Sciences of the United States of America, 117(42), 26020–26030. doi: 10.1073/pnas.2008303117

Hantschel, O., Nagar, B., Guettler, S., Kretzschmar, J., Dorey, K., Kuriyan, J., & Superti-Furga, G. (2003). A myristoyl/phosphotyrosine switch regulates c-Abl. Cell, 112(6), 845–857. doi: 10.1016/s0092-8674(03)00191-0

Hashimoto, S., Iwamatsu, A., Ishiai, M., Okawa, K., Yamadori, T., Matsushita, M., … Tsukada, S. (1999). Identification of the SH2 domain binding protein of Bruton’s tyrosine kinase as BLNK--functional significance of Btk-SH2 domain in B-cell antigen receptor-coupled calcium signaling. Blood, 94(7), 2357–2364.

Hein, M. Y., Hubner, N. C., Poser, I., Cox, J., Nagaraj, N., Toyoda, Y., … Mann, M. (2015). A human interactome in three quantitative dimensions organized by stoichiometries and abundances. Cell, 163(3), 712–723. doi: 10.1016/j.cell.2015.09.053

Hendriks, R. W., Yuvaraj, S., & Kil, L. P. (2014). Targeting Bruton’s tyrosine kinase in B cell malignancies. Nature Reviews. Cancer, 14(4), 219–232. doi: 10.1038/nrc3702

Huang, W. Y. C., Alvarez, S., Kondo, Y., Lee, Y. K., Chung, J. K., Lam, H. Y. M., … Groves, J. T. (2019). A molecular assembly phase transition and kinetic proofreading modulate Ras activation by SOS. Science, 363(6431), 1098–1103. doi: 10.1126/science.aau5721

Huang, W. Y. C., Chiang, H.-K., & Groves, J. T. (2017). Dynamic Scaling Analysis of Molecular Motion within the LAT:Grb2:SOS Protein Network on Membranes. Biophysical Journal, 113(8), 1807–1813. doi: 10.1016/j.bpj.2017.08.024

Huang, W. Y. C., Ditlev, J. A., Chiang, H.-K., Rosen, M. K., & Groves, J. T. (2017). Allosteric modulation of grb2 recruitment to the intrinsically disordered scaffold protein, LAT, by remote site phosphorylation. Journal of the American Chemical Society, 139(49), 18009–18015. doi: 10.1021/jacs.7b09387

Huang, W. Y. C., Yan, Q., Lin, W.-C., Chung, J. K., Hansen, S. D., Christensen, S. M., … Groves, J. T. (2016). Phosphotyrosine-mediated LAT assembly on membranes drives kinetic bifurcation in recruitment dynamics of the Ras activator SOS. Proceedings of the National Academy of Sciences of the United States of America, 113(29), 8218–8223. doi: 10.1073/pnas.1602602113

Hyvönen, M., & Saraste, M. (1997). Structure of the PH domain and Btk motif from Bruton’s tyrosine kinase: molecular explanations for X-linked agammaglobulinaemia. The EMBO Journal, 16(12), 3396–3404. doi: 10.1093/emboj/16.12.3396

Joseph, R. E., Wales, T. E., Fulton, D. B., Engen, J. R., & Andreotti, A. H. (2017). Achieving a graded immune response: BTK adopts a range of active/inactive conformations dictated by multiple interdomain contacts. Structure, 25(10), 1481–1494.e4. doi: 10.1016/j.str.2017.07.014

Kaizuka, Y., & Groves, J. T. (2004). Structure and dynamics of supported intermembrane junctions. Biophysical Journal, 86(2), 905–912. doi: 10.1016/S0006-3495(04)74166-1

Kim, W., Kim, E., Min, H., Kim, M. G., Eisenbeis, V. B., Dutta, A. K., … Seong, R. H. (2019). Inositol polyphosphates promote T cell-independent humoral immunity via the regulation of Bruton’s tyrosine kinase. Proceedings of the National Academy of Sciences of the United States of America, 116(26), 12952–12957. doi: 10.1073/pnas.1821552116

Knight, J. D., & Falke, J. J. (2009). Single-molecule fluorescence studies of a PH domain: new insights into the membrane docking reaction. Biophysical Journal, 96(2), 566–582. doi: 10.1016/j.bpj.2008.10.020

Koretzky, G. A., Abtahian, F., & Silverman, M. A. (2006). SLP76 and SLP65: complex regulation of signalling in lymphocytes and beyond. Nature Reviews. Immunology, 6(1), 67–78. doi: 10.1038/nri1750

Kurosaki, T., & Tsukada, S. (2000). BLNK: connecting Syk and Btk to calcium signals. Immunity, 12(1), 1–5. doi: 10.1016/s1074-7613(00)80153-3

Lander, G. C., Estrin, E., Matyskiela, M. E., Bashore, C., Nogales, E., & Martin, A. (2012). Complete subunit architecture of the proteasome regulatory particle. Nature, 482(7384), 186–191. doi: 10.1038/nature10774

Lin, C.-C., Melo, F. A., Ghosh, R., Suen, K. M., Stagg, L. J., Kirkpatrick, J., … Ladbury, J. E. (2012). Inhibition of basal FGF receptor signaling by dimeric Grb2. Cell, 149(7), 1514–1524. doi: 10.1016/j.cell.2012.04.033

Lin, C.-W., Nocka, L. M., Stinger, B., DeGrandchamp, J., Lew, N., Alvarez, S., … Groves, J. T. (2021). A two-component protein condensate of EGFR and Grb2 regulates Ras activation at the membrane. BioRxiv. doi: 10.1101/2021.12.12.472247

Lin, C.-W., Nocka, L. M., Stinger, B. L., DeGrandchamp, J. B., Lew, L. J. N., Alvarez, S., … Groves, J. T. (2022). A two-component protein condensate of the EGFR cytoplasmic tail and Grb2 regulates Ras activation by SOS at the membrane. Proceedings of the National Academy of Sciences of the United States of America, 119(19), e2122531119. doi: 10.1073/pnas.2122531119

Lin, J. J., O’Donoghue, G. P., Wilhelm, K. B., Coyle, M. P., Low-Nam, S. T., Fay, N. C., … Groves, J. T. (2020). Membrane Association Transforms an Inert Anti-TCRβ Fab’ Ligand into a Potent T Cell Receptor Agonist. Biophysical Journal, 118(12), 2879–2893. doi: 10.1016/j.bpj.2020.04.018

Liu, S. K., Fang, N., Koretzky, G. A., & McGlade, C. J. (1999). The hematopoietic-specific adaptor protein gads functions in T-cell signaling via interactions with the SLP-76 and LAT adaptors. Current Biology, 9(2), 67–75. doi: 10.1016/s0960-9822(99)80017-7

Lowenstein, E. J., Daly, R. J., Batzer, A. G., Li, W., Margolis, B., Lammers, R., … Schlessinger, J. (1992). The SH2 and SH3 domain-containing protein GRB2 links receptor tyrosine kinases to ras signaling. Cell, 70(3), 431–442. doi: 10.1016/0092-8674(92)90167-b

Maignan, S., Guilloteau, J. P., Fromage, N., Arnoux, B., Becquart, J., & Ducruix, A. (1995). Crystal structure of the mammalian Grb2 adaptor. Science, 268(5208), 291–293. doi: 10.1126/science.7716522

Mayer, B. J., Jackson, P. K., Van Etten, R. A., & Baltimore, D. (1992). Point mutations in the abl SH2 domain coordinately impair phosphotyrosine binding in vitro and transforming activity in vivo. Molecular and Cellular Biology, 12(2), 609–618. doi: 10.1128/mcb.12.2.609-618.1992

Moarefi, I., LaFevre-Bernt, M., Sicheri, F., Huse, M., Lee, C. H., Kuriyan, J., & Miller, W. T. (1997). Activation of the Src-family tyrosine kinase Hck by SH3 domain displacement. Nature, 385(6617), 650–653. doi: 10.1038/385650a0

Moroco, J. A., Craigo, J. K., Iacob, R. E., Wales, T. E., Engen, J. R., & Smithgall, T. E. (2014). Differential sensitivity of Src-family kinases to activation by SH3 domain displacement. Plos One, 9(8), e105629. doi: 10.1371/journal.pone.0105629

Nagar, B., Hantschel, O., Young, M. A., Scheffzek, K., Veach, D., Bornmann, W., … Kuriyan, J. (2003). Structural basis for the autoinhibition of c-Abl tyrosine kinase. Cell, 112(6), 859–871. doi: 10.1016/s0092-8674(03)00194-6

Nye, J. A., & Groves, J. T. (2008). Kinetic control of histidine-tagged protein surface density on supported lipid bilayers. Langmuir: The ACS Journal of Surfaces and Colloids, 24(8), 4145–4149. doi: 10.1021/la703788h

Olivier, J. P., Raabe, T., Henkemeyer, M., Dickson, B., Mbamalu, G., Margolis, B., … Pawson, T. (1993). A Drosophila SH2-SH3 adaptor protein implicated in coupling the sevenless tyrosine kinase to an activator of Ras guanine nucleotide exchange, Sos. Cell, 73(1), 179–191. doi: 10.1016/0092-8674(93)90170-u

Ozdener, F., Dangelmaier, C., Ashby, B., Kunapuli, S. P., & Daniel, J. L. (2002). Activation of phospholipase Cgamma2 by tyrosine phosphorylation. Molecular Pharmacology, 62(3), 672–679. doi: 10.1124/mol.62.3.672

Reif, K., Buday, L., Downward, J., & Cantrell, D. A. (1994). SH3 domains of the adapter molecule Grb2 complex with two proteins in T cells: the guanine nucleotide exchange protein Sos and a 75-kDa protein that is a substrate for T cell antigen receptor-activated tyrosine kinases. The Journal of Biological Chemistry, 269(19), 14081–14087.

Rip, J., Van Der Ploeg, E. K., Hendriks, R. W., & Corneth, O. B. J. (2018). The role of bruton’s tyrosine kinase in immune cell signaling and systemic autoimmunity. Critical Reviews in Immunology, 38(1), 17–62. doi: 10.1615/CritRevImmunol.2018025184

Rodriguez, R., Matsuda, M., Perisic, O., Bravo, J., Paul, A., Jones, N. P., … Katan, M. (2001). Tyrosine residues in phospholipase Cgamma 2 essential for the enzyme function in B-cell signaling. The Journal of Biological Chemistry, 276(51), 47982–47992. doi: 10.1074/jbc.M107577200

Scharenberg, A. M., Humphries, L. A., & Rawlings, D. J. (2007). Calcium signalling and cell-fate choice in B cells. Nature Reviews. Immunology, 7(10), 778–789. doi: 10.1038/nri2172

Seeliger, M. A., Young, M., Henderson, M. N., Pellicena, P., King, D. S., Falick, A. M., & Kuriyan, J. (2005). High yield bacterial expression of active c-Abl and c-Src tyrosine kinases. Protein Science, 14(12), 3135–3139. doi: 10.1110/ps.051750905

Shah, N. H., Amacher, J. F., Nocka, L. M., & Kuriyan, J. (2018). The Src module: an ancient scaffold in the evolution of cytoplasmic tyrosine kinases. Critical Reviews in Biochemistry and Molecular Biology, 53(5), 535–563. doi: 10.1080/10409238.2018.1495173

Shi, T., Niepel, M., McDermott, J. E., Gao, Y., Nicora, C. D., Chrisler, W. B., … Wiley, H. S. (2016). Conservation of protein abundance patterns reveals the regulatory architecture of the EGFR-MAPK pathway. Science Signaling, 9(436), rs6. doi: 10.1126/scisignal.aaf0891

Su, X., Ditlev, J. A., Hui, E., Xing, W., Banjade, S., Okrut, J., … Vale, R. D. (2016). Phase separation of signaling molecules promotes T cell receptor signal transduction. Science, 352(6285), 595–599. doi: 10.1126/science.aad9964

Tsukada, S., Saffran, D. C., Rawlings, D. J., Parolini, O., Allen, R. C., Klisak, I., … Quan, S. (1993). Deficient expression of a B cell cytoplasmic tyrosine kinase in human X-linked agammaglobulinemia. Cell, 72(2), 279–290. doi: 10.1016/0092-8674(93)90667-f

Wang, Q., Pechersky, Y., Sagawa, S., Pan, A. C., & Shaw, D. E. (2019). Structural mechanism for Bruton’s tyrosine kinase activation at the cell membrane. Proceedings of the National Academy of Sciences of the United States of America, 116(19), 9390–9399. doi: 10.1073/pnas.1819301116

Wang, Q., Vogan, E. M., Nocka, L. M., Rosen, C. E., Zorn, J. A., Harrison, S. C., & Kuriyan, J. (2015). Autoinhibition of Bruton’s tyrosine kinase (Btk) and activation by soluble inositol hexakisphosphate. ELife, 4, e06074. doi: 10.7554/eLife.06074

Weiss, A., & Littman, D. R. (1994). Signal transduction by lymphocyte antigen receptors. Cell, 76(2), 263–274. doi: 10.1016/0092-8674(94)90334-4

Yeung, W., Kwon, A., Taujale, R., Bunn, C., Venkat, A., & Kannan, N. (2021). Evolution of functional diversity in the holozoan tyrosine kinome. Molecular Biology and Evolution, 38(12), 5625–5639. doi: 10.1093/molbev/msab272

